# The complex origins of strigolactone signalling in land plants

**DOI:** 10.1101/102715

**Authors:** Rohan Bythell-Douglas, Carl J. Rothfels, Dennis W.D. Stevenson, Gane Ka-Shu Wong, David C. Nelson, Tom Bennett

**Author notes:** corresponding author: Tom Bennett.

## Abstract

Strigolactones (SLs) are a class of plant hormones that control many aspects of plant growth. The SL signalling mechanism is homologous to that of karrikins (KARs), smoke-derived compounds that stimulate seed germination. In angiosperms, the SL receptor is an α/β hydrolase known as DWARF14 (D14); its close homologue, KARRIKIN INSENSITIVE2 (KAI2), functions as a KAR receptor, and likely recognizes an uncharacterized, endogenous signal. Previous phylogenetic analyses have suggested that the KAI2 lineage is ancestral in land plants, and that canonical D14-type SL receptors only arose in seed plants; this is paradoxical, however, as non-vascular plants synthesize and respond to SLs. Here, we have used a combination of phylogenetic and structural approaches to re-assess the evolution of the D14/KAI2 family in land plants. We analyzed 339 members of the D14/KAI2 family from land plants and charophyte algae. Our phylogenetic analyses show that the divergence between the eu-KAI2 lineage and the DDK (<u>D</u>14/<u>D</u>LK2/<u>K</u>AI2) lineage that includes D14 occurred very early in land plant evolution. We identify characteristic structural features of D14 and KAI2 proteins, and use homology modelling to show that the earliest members of the DDK lineage structurally resemble KAI2, and not D14 proteins. Furthermore, we show that probable SL receptors in non-seed plants do not have D14-like structure. Our results suggest that SL perception has relatively relaxed structural requirements, and that the evolution from KAI2-like to D14-like protein structure in the DDK lineage may have been driven by interactions with protein partners, rather than being required for SL perception itself.

## INTRODUCTION

Plant hormones are a key link between environmental stimuli and development, allowing local information to be used systemically across the plant body. Strigolactones (SLs) are a recently identified class of terpenoid lactone hormone that neatly epitomise this concept. SLs are primarily synthesised by a core pathway involving a carotene isomerase (DWARF27), two carotenoid cleavage dioxygenases (CCD7 and CCD8)(Al-Babili et al, 2015), and a cytochrome P450 enzyme (MAX1). SL synthesis is strongly upregulated by phosphate deficiency in the rhizosphere (Lopez-Raez et al, 2008), increasing the pool of SL molecules in the root. In many flowering plants (angiosperms), SLs are exuded into the soil through the action of specific SL transporters, and serve to attract mycorrhizal fungi (Borghi et al, 2016); the resulting symbioses provide the plants with phosphate in exchange for reduced carbon. SLs also act locally to regulate root system architecture; the precise effects seem to vary from species to species, but increased SL levels may promote increased nutrient foraging (Mattys et al, 2016). Finally, a significant proportion of the SL pool produced in the root is transported into the shoot system via the xylem (Kohlen et al, 2011), where it has a well-defined set of effects on shoot growth and development (Smith & Waters, 2012; Waters et al, 2017). SL has inhibitory effect on shoot branching, thereby coupling shoot growth to nutrient availability (Kohlen et al, 2011). SL responses thus form an integrated stimulus-response system acting over long distances both within the plant body and its immediate environment.

Like several other plant hormonal signalling pathways, canonical SL signalling is mediated through ubiquitin-mediated degradation of target proteins (reviewed in Waters et al, 2017). The SL receptors for this signalling pathway are members of the DWARF14 (D14) class of α/β hydrolase proteins, which are an unusual combination of enzyme and receptor (de Saint Germain et al, 2016; Yao et al, 2016). D14 proteins bind and then cleave SL molecules, producing an intermediate molecule (CLIM) that is covalently bound to the receptor (de Saint Germain et al, 2016; Yao et al, 2016). SL signalling is mediated through the interaction of D14 with the MORE AXILLARY GROWTH2 (MAX2) class of F-box protein, which form part of an SCF (SKP1-CULLIN-F-BOX) E3-ubiquitin ligase (Stirnberg et al, 2002; Stirnberg et al, 2007; Hamiaux et al, 2012; Zhao et al, 2015). Together, the covalent binding of CLIM and the interaction with SCF^MAX2^ allow D14 to undergo a stable conformational change that drives onward signalling (de Saint Germain et al, 2016; Yao et al, 2016). Although other targets have been proposed (Nakamura et al, 2013; Wang et al, 2013), it is now clear that the principal proteolytic targets of SL signalling are proteins of the SMAX1-LIKE7/DWARF53 (SMXL7/D53) class (Zhou et al, 2013; Jiang et al, 2013; Soundappan et al, 2015; Wang et al, 2015; Liang et al, 2016; Bennett et al, 2016). The exact sequence of events is unclear, but it is probably after conformational change that D14 stably recruits SMXL7 to the complex; certainly, the D14-SMXL7 interaction is enhanced by SL (Zhou et al, 2013; Jiang et al, 2013; Wang et al, 2015; Liang et al, 2016). Events downstream of SMXL7 degradation are currently poorly defined; SMXL7 has been proposed to act both transcriptionally and non-transcriptionally (Bennett & Leyser, 2014; Waters et al, 2017). It may be that SMXL7 is a multi-functional protein that can regulate multiple cellular processes (Liang et al, 2016).

Intriguingly, a second pathway in angiosperms signals through SCF^MAX2^, forming a biochemical and evolutionary parallel to SL signalling. This pathway is defined by the KARRIKIN INSENSITIVE2 (KAI2) α/β hydrolase protein, a close relative of D14. *kai2* mutants have a range of developmental phenotypes (Waters et al, 2012; Soundappan et al, 2015; Bennett et al, 2016), and are insensitive to the germination-promoting effects of smoke-derived ‘karrikins’ (KARs)(Waters et al, 2012). It has been hypothesized that karrikins promote germination by mimicking an as-yet-unidentified endogenous KAI2 ligand (‘KL’)(Flematti et al, 2013; Conn & Nelson, 2016). MAX2 is required for both responses to karrikins and for other aspects of KAI2-dependent signalling (Nelson et al, 2011; Soundappan et al, 2015; Bennett et al, 2016). Furthermore, the presumptive proteolytic targets of KAI2-SCF^MAX2^ signalling are close homologues of SMXL7; in Arabidopsis, these are SMXL2 and SMAX1 (SUPPRESSOR OF MAX2 1) itself. Mutation of *SMAX1* and *SMXL2* suppresses the *kai2*-related phenotypes present in the *max2* mutant, producing phenotypes that mimic constitutive karrikin responses (Stanga et al, 2013; Soundappan et al, 2015; Stanga et al, 2016). In the Arabidopsis genome, there are further homologues of *D14* and *SMAX1*, namely *DWARF14-LIKE2* (*DLK2*) and *SMXL3, SMXL4* and *SMXL5*, but the function of these proteins and their relationship to SL/ KL signalling is currently unclear (Waters et al, 2012; Stanga et al, 2013).

The evolutionary history of SLs represents an intriguing and unresolved problem. SLs have been identified in most land plant groups, and in some related groups of charophyte algae (Delaux et al, 2012). However, unambiguous *CCD8* orthologues have not been identified in charophytes or liverworts (a possible sister group to other land plants)(Wickett et al, 2014; Delaux et al, 2012). Moreover, *ccd8* mutants in the moss *Physcomitrella patens* still produce some SLs (Proust et al, 2011), which suggests there may be alternative pathways for SL synthesis (Waldie et al, 2014; Waters et al, 2017). Even more uncertainty surrounds the origin of the canonical SL signalling pathway. Unambiguous *D14* orthologues have only been identified in seed plants (gymnosperms and angiosperms), and seem to be absent from mosses and liverworts (Delaux et al, 2012; Waters et al, 2015). Conversely, it has been suggested that unambiguous *KAI2* orthologues are present in charophytes, liverworts and mosses (Delaux et al, 2012). This has led to the suggestion that KAI2 proteins could function as receptor for SLs in non-vascular plants, or that SL signalling occurs by non-canonical mechanisms in these lineages (Bennett & Leyser, 2014; Waters et al, 2017). Supporting the plausibility of the former hypothesis, it was recently shown that SL receptors evolved from *KAI2* paralogs in parasitic plants within the Orobanchaceae (Conn et al., 2015; Tsuchiya et al., 2015; Toh et al., 2015). In addition, *MAX2* orthologues have so far only been identified in land plants (Challis et al, 2013), and while *MAX2* is present in *P. patens*, *Ppmax2* mutants do not resemble *Ppccd8* mutants, suggesting MAX2 may not be involved in SL signalling in mosses (de Saint Germain et al, 2013; Bennett & Leyser, 2014). Thus, even if KAI2 proteins can act as SL receptors in mosses, they may not act through SCF^MAX2^–mediated protein degradation. SMXL proteins are present in *P. patens* but their function has not been investigated. Thus, while there is clear evidence for SL sensitivity in mosses, it is possible this occurs through separate mechanisms to those in angiosperms. This would contrast strongly with the auxin signalling pathway for instance, which is completely conserved throughout land plants (Lavy et al, 2016; Flores-Sandoval et al, 2015; Kato et al, 2015).

To resolve the evolutionary history of SL signalling, we have undertaken a major phylogenetic re-assessment of the *D14*/*KAI2* family. We identified 339 *D14/KAI2* homologues, sampled from each major land plant group using completed genome sequences and transcriptome assemblies from the 1KP project. We reconstructed the evolution of the family using a range of methods, and these analyses converged on a common topology. Our results suggest a deep duplication in the *D14*/*KAI2* family at the base of land plants, leading to a ‘*eu-KAI2*’ clade and a ‘*DDK’* (*<u>D</u>14*-*<u>D</u>LK2*-*<u>K</u>AI2*) clade that contains the characterized angiosperm D14 SL receptors. We analysed the primary structure of these proteins, and found that members of eu-KAI2 clade contain highly conserved features. Conversely, as a whole, the DDK clade has weak structural conservation, though individual clades within the family are well conserved. We identified sets of residues that define the D14 and KAI2 receptors, and use these to assess the nature of other DDK proteins. Finally, we used homology modelling of proteins in the DDK lineage to assess their possible ligand binding capabilities. Our results suggest that the lineage leading to D14 proteins arose early in land plant evolution, and that highly specialized D14 SL receptors in gymnosperms gradually evolved from KAI2-like proteins. We also identify several groups of divergent proteins that appear to have lost the conserved residues that comprise the MAX2-interface, and may represent novel receptor proteins.

## RESULTS

### Preliminary analysis of the *D14/KAI2* family

In order to understand the evolution of the *D14*/*KAI2* family with greater resolution, we obtained 339 sequences from 143 species, representing the major lineages of land plants and charophyte algae (Table 1). All preliminary phylogenetic analyses placed *D14*/*KAI2* family members into unambiguous taxon-specific clades such as angiosperm *KAI2* or gymnosperm *D14* (Table 2). Understanding the interrelationship of these taxon-level clades therefore seemed to be key to understanding the evolution of the *D14*/*KAI2* family. Sequences from each major land plant taxon grouped into at least two distinct clades, except for the hornworts, in which all sequences grouped into a single clade (Table 2).

**Table 1:**
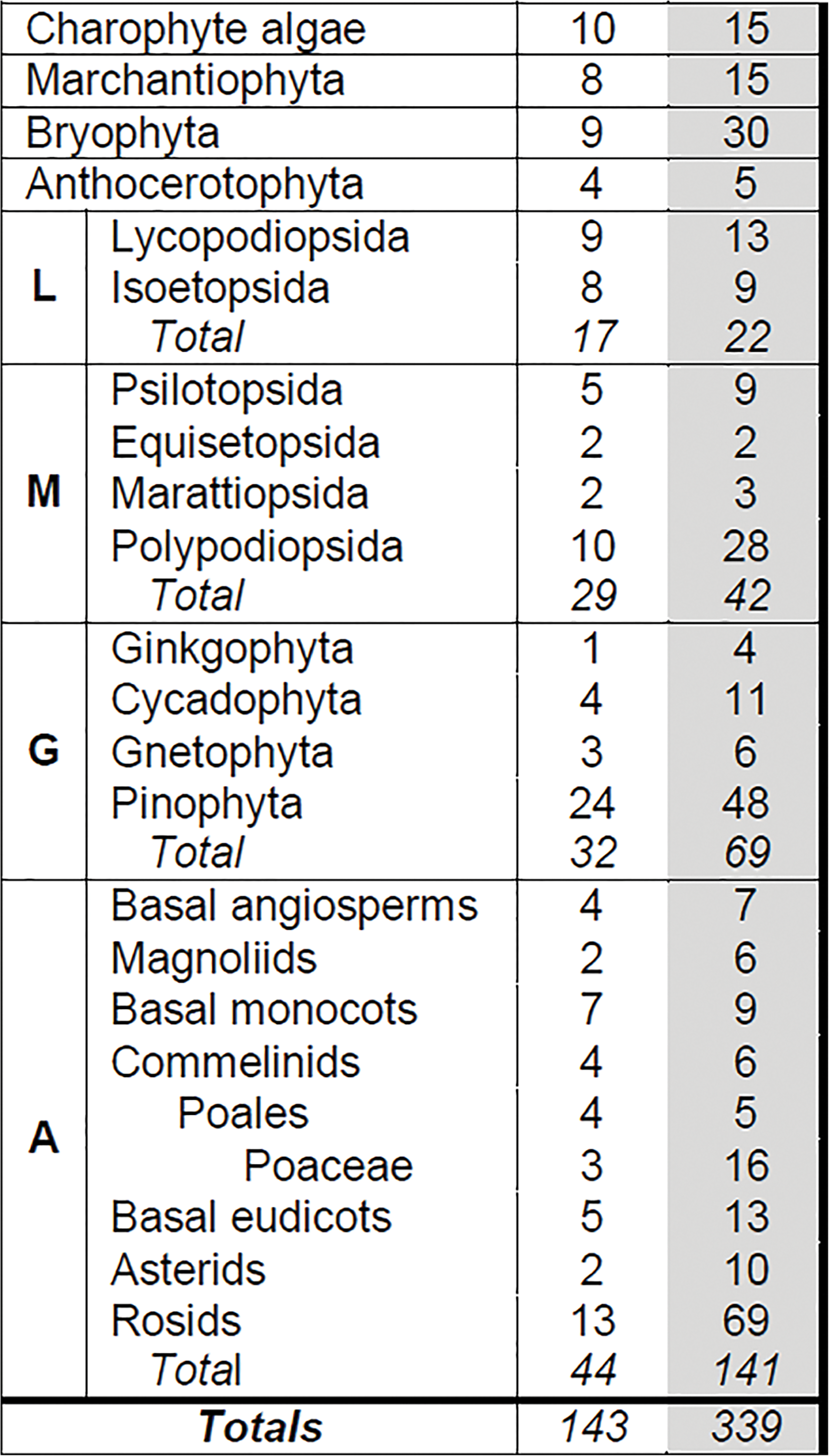
Sampling of *D14/KAI2* family members. Table showing *D14*/*KAI2* family sampling rates across the plant kingdom. The primary taxonomic divisions are shown at the left; lycophytes (L), monilophytes (M), gymnosperms (G) and angiosperms (A) are further broken down into major subgroups. The number of species (unshaded) and the number of sequences (shaded) obtained from each taxon are shown. Numbers for the Poaceae are shown separately from other Poales, which are in turn shown separately from other commelinids.

**Table 2:**
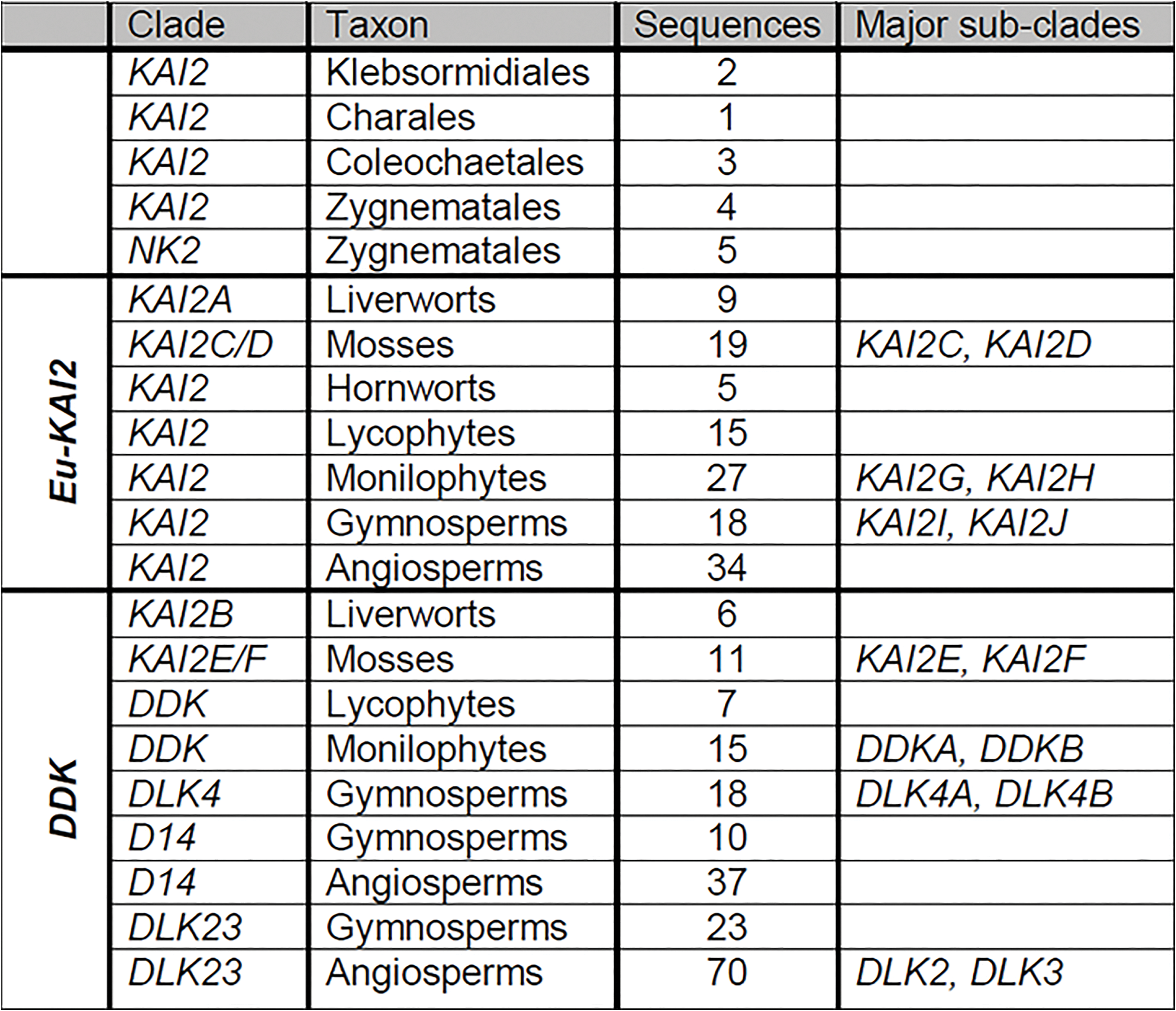
Major clades in the *D14/KAI2* family. Table showing major clades in the *D14/KAI2* family, as defined at the level of major taxonomic groups. Almost all sequences in the family unambiguously group into one of these clades. Within some clades there are major sub-clades where the lineage has been duplicated; these are listed at the right. Our analysis suggests that land plant *D14*/*KAI2* proteins group into two super-clades, *eu-KAI2* and *DDK*, as indicated on the left of the table.

From species in the charophyte orders Klebsormidiales, Charales, and Coleochaetales we only obtained a single sequence per genome, all of which superficially resembled *KAI2*. However, from several species in the Zygnematales we obtained two distinct types of sequences, one resembling *KAI2* and the other not, which we named *NOT KAI2* (*NK2*). In recent analyses the Zygnematales have been identified as a good candidates for the sister group to land plants, even though morphological analyses have traditionally favoured the Charales in this respect (Wodniok et al, 2011; Timme et al, 2012; Doyle et al, 2013; Wickett et al, 2014). If this reconstruction is correct, the two lineages present in Zygnematales could be evidence that the duplication in the *D14*/*KAI2* family occurred before the land plant-Zygnematales split. However, in our analyses *NK2* sequences grouped with other charophyte *KAI2* sequences (Figure 1), and they have highly divergent characteristics, unlike any other members of the *D14*/*KAI2* family. We therefore propose that these genes represent a Zygnematales-specific duplication.

**Figure 1:**
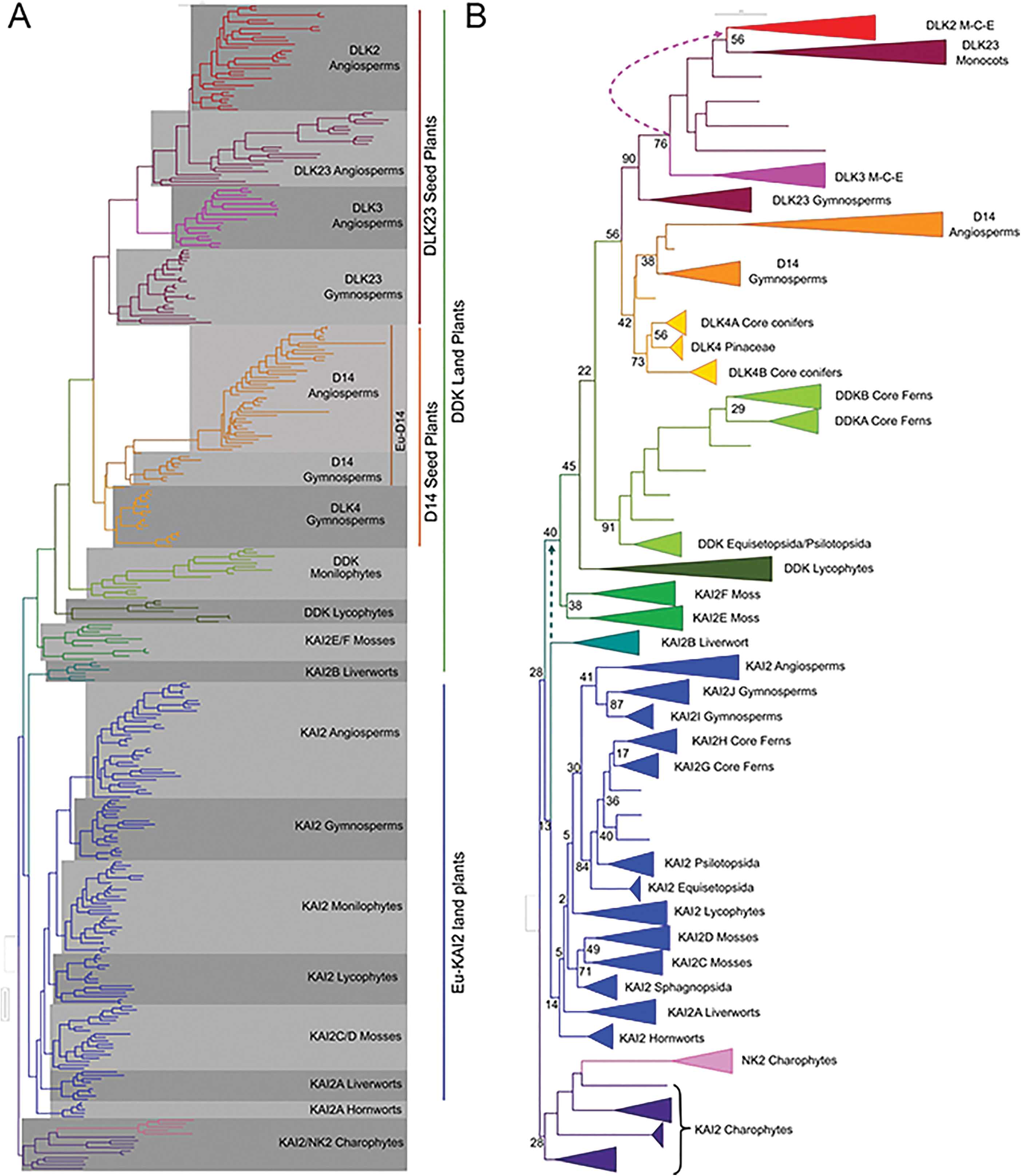
The *eu-KAI2* and *DDK* super-clades diverged early in land plant evolution. Codon-level phylogenetic analysis implemented in GARLI on the whole *D14*/*KAI2* family (339 sequences from 143 species). This analysis was performed using an optimized character set (see materials and methods). Trees were rooted with charophyte sequences, consistent with contemporary notions of plant organismal phylogeny. Dotted lines indicate alternative positions for the indicated clades that would increase the parsimony of the tree. **A)** Phylogram showing the ‘most likely’ tree from GARLI analysis, labelled to show the high-order relationships between the major clades (as described in Table 2). **B)** Cladogram depicting the phylogenetic tree from A) in simplified form. Major clades and sub-clades (as listed in Table 2) are collapsed. Numbers associated with internal branches denote maximum likelihood bootstrap support (% support). M-C-E = magnoliids-chloranthales-eudicots

### Multiple analyses support an early origin for the DDK super-clade

To explore the evolution of the *D14*/*KAI2* family, we performed maximum likelihood phylogenetic analyses using both nucleotide and amino acid sequence data, implemented in PhyML and GARLI (Guindon et al, 2010; Zwickl et al, 2006). Preliminary analyses were run on a ‘maximum’ alignment of 780 nucleotides from all 339 sequences, and the resulting trees rooted with charophyte sequences. However, we found that lycophyte *KAI2* sequences (particularly those from Selaginella spp.) tended to be misplaced near the root of the tree. This is a recognized problem in land plant phylogenies, caused by divergent codon usage in lycophytes (particularly Selaginella), which resembles that of charophytes (Cox et al, 2014). We were able to improve the overall tree topology, resulting in more realistic branching orders, by using progressively smaller and more conservative alignments (Figure 1, Supplementary Figure 1A). If we removed the charophyte and lycophyte sequences (leaving 296 sequences from 122 species), we were able to recover the same basic topology, but using the maximum DNA alignment (Figure 2, Supplementary Figure 1B).

**Figure 2:**
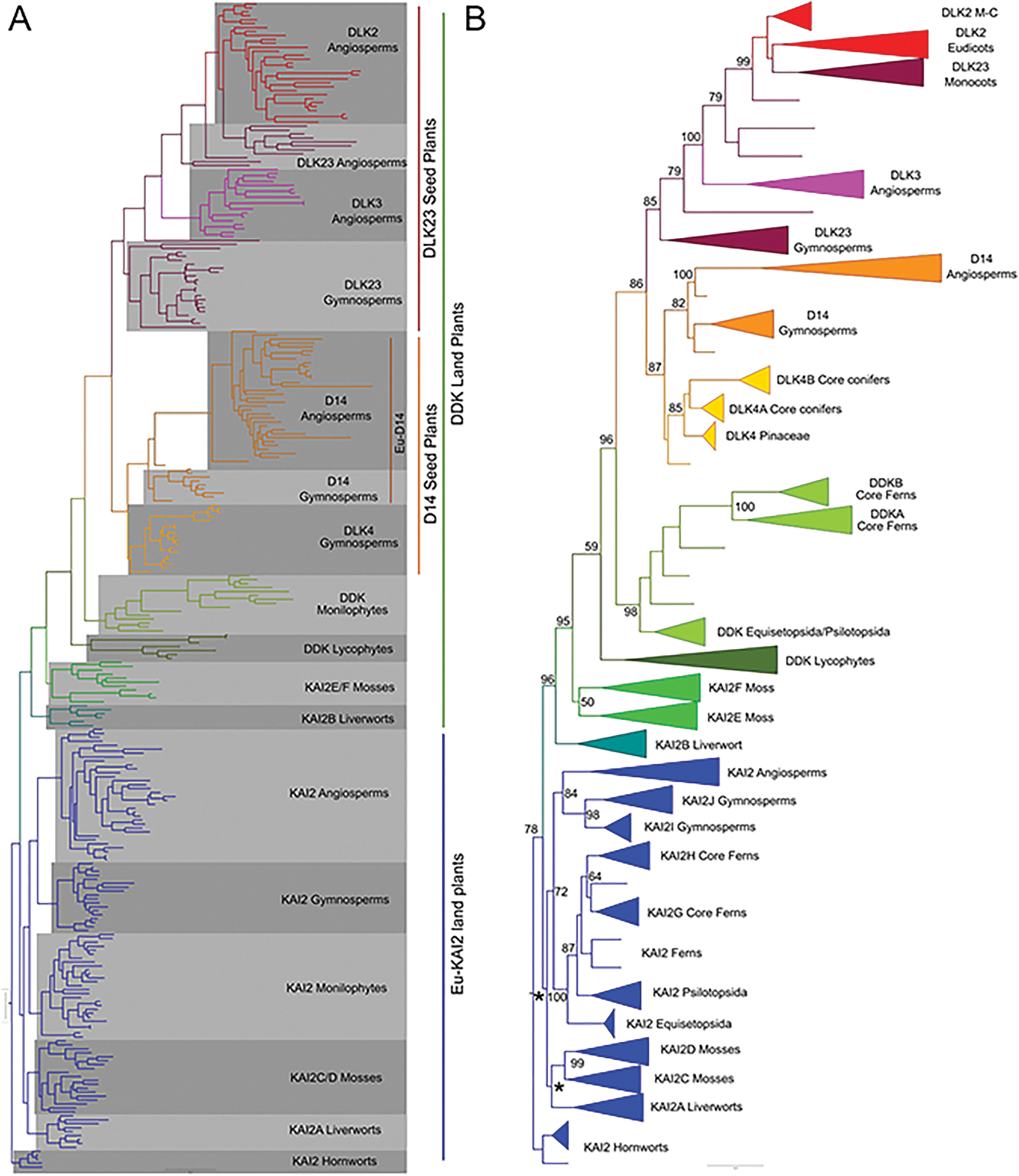
The *eu-KAI2* and *DDK* super-clades diverged early in land plant evolution. Nucleotide-level phylogenetic analysis implemented in GARLI on the *D14*/*KAI2* family, minus charophyte and lycophyte *KAI2* sequences (296 sequences). Trees were rooted with hornwort *KAI2* sequences by comparison with Figure 1. This analysis was performed using the full-length dataset (780 characters). **A)** Phylogram showing the ‘most likely’ tree from GARLI analysis, labelled to show the high-order relationships between the major clades (as described in Table 2). **B)** Cladogram depicting the phylogenetic tree from A) in simplified form. Major clades and sub-clades (as listed in Table 2) are collapsed. Numbers associated with internal branches denote maximum likelihood bootstrap support (% support); values below 50 are indicated with an asterisk *. M-C = magnoliids/chloranthales

Irrespective of the underlying alignment and methodology, all analyses agreed on a basic topology for the family, with a deep duplication near the base of the land plants creating two super-clades. The first lineage contains *KAI2* sequences from angiosperms and closely related sequences from gymnosperms, monilophytes, lycophytes, mosses, and liverworts; we therefore named this clade *eu-KAI2* (Table 2). The second super-clade contains sequences from mosses that have previously been described as *KAI2*-like (Waters et al, 2012; Waters et al, 2015; Lopez-Obando et al, 2016), sequences from lycophytes and monilophytes that do not resemble known proteins, the previously characterized *D14* and *DLK2* genes from angiosperms, and homologous genes from gymnosperms (Table 2). To reflect the mixed composition of this clade, we named it ‘*DDK*’ (for *D14*/*DLK2*/*KAI2*); we also used this name for the monilophyte and lycophyte sequences in the clade. The lycophyte *DDK* group contains the *Selaginella moellendorffi* gene previously described as ‘*SmKAI2b*’ (Waters et al, 2015), but we believe *DDK* designation better reflects the evolutionary context of these proteins. We observed some variation in the composition of the *eu-KAI2* clade, partly as a result of the erratic behavior of the lycophyte *KAI2* sequences. However, the moss *KAI2E*/*F*, lycophyte *DDK*, monilophyte *DDK*, gymnosperm *D14*, *DLK4*, *DLK23* and angiosperm *D14* and *DLK23* clades were associated into a single large clade in every analysis we performed, though the internal branching order did vary somewhat between analyses. This basic topology was evident even in very early analyses (Supplementary Data 1E).

Only two clades were inconsistently placed. The hornwort *KAI2* clade is the most problematic in our analyses, mirroring the uncertainty about the position of the hornworts themselves in organismal phylogeny (Wickett et al, 2014). In some analyses the hornwort *KAI2* clade is placed in the *eu-KAI2* lineage, between mosses and lycophytes (Supplementary Data 1C). Alternatively, it is also placed at the base of the *eu-KAI2* lineage (Figure 1), or as a sister clade to all other land plant *D14*/*KAI2* sequences (Figure 2, Supplementary Data 1A). None of these positions alters the interpretation of a deep duplication in the family, but they do affect its inferred timing. The liverwort *KAI2B* clade occurs either at the base of the *DDK* or *eu-KAI2* lineages in different trees. In analyses performed without charophyte and lycophyte *KAI2* sequences it is always associated with the *DDK* lineage (Figure 2, Supplementary Data 1B,C). This is also the case in some analyses including charophyte sequences (Supplementary Data 1A). The position at the base of the *eu-KAI2* clade in some trees is likely to be erroneous, and probably caused by the slight misplacement of charophyte sequences. For instance, the liverwort-hornwort-liverwort branching order at the base of the *eu-KAI2* clade in Figure 1 is highly improbable. Rooting this tree with the hornwort *KAI2* clade (to match Figure 2) produces balanced *eu-KAI2* and *DDK* clades, with realistic branching order, except for the inclusion of the charophyte sequences as an in-group within the *DDK* clade (Supplementary Data 1E). We believe the most parsimonious scenario is that *KAI2B* is part of the *DDK* clade.

Collectively, our phylogenetic analyses push the origin of the *D14* lineage back much earlier than proposed in previous phylogenies that suggested an origin in the vascular plants (Waters et al, 2012) or within the seed plants (Waters et al, 2015). They resolve the enigmatic placement of *SmKAI2b* and divergent *KAI2* sequences from *P. patens* in previous phylogenies (Waters et al, 2015; Lopez-Obando et al, 2015). They also provide a convincing explanation for the presence of two distinct *D14*/*KAI2* clades in most major plant groups. Key to this reconstruction topology is the placement of liverwort and moss clades with apparently KAI2-like primary protein structure (KAI2B and KAI2E/F respectively) in the *DDK* lineage. We wanted to test the robustness of this somewhat unexpected conclusion, and used a variety of methods to do so.

Non-parametric bootstrap analyses performed in GARLI did not provide very high levels of support for most of the nodes along the backbone of the tree (Figure 1). However, bootstrap values were higher in reconstructions that excluded charophyte and lycophyte *KAI2* sequences (Figure 2). We next tested whether the recovered topology was stable to perturbations in the dataset. We re-ran our analysis multiple times, removing each *DDK* clade in turn (see Materials and Methods). Our analysis suggests that the placement of *KAI2B* is sensitive to the dataset used, but that the rest of the *DDK* clade is very stably associated (Supplementary Data 1D). Finally, we assessed whether our general topology is congruent with previous analyses. We observed that in Waters et al (2012), the *Marchantia polymorpha KAI2A* and *KAI2B* sequences do not group together, and neither do the *P. patens KAI2C*/*D* and *KAI2E*/*F*. This is consistent with our analyses. We repeated our analysis using a set of sequences pruned to match Waters et al (2012) and found essentially the same tree as in that study (Supplementary Data 1F). Furthermore, if we rooted the tree with a *eu-KAI2* sequence, we observed essentially the same topology as in our study (Supplementary Data 1F). This shows that the difference in final topology between our study and Waters et al (2012) does not result from any particular methodological differences, but from our more densely populated sequence set.

### Diverse evolutionary histories in the *D14/KAI2* family

From our phylogenetic reconstruction, it is apparent that the two super-clades of the *D14*/*KAI2* family appear to have rather different evolutionary trajectories (Figure 3). Within the *eu-KAI2* super-clade there is a single clade for each major plant group (e.g. angiosperm *KAI2*). Within these taxon-specific clades, there have apparently been some early duplications. For instance, *KAI2C* and *KAI2D* clades are widely represented among extant mosses, although not in the Sphagnopsida, suggesting the duplication occurred after the separation of the Sphagnopsida and other mosses (Figure 1). Similarly, the separation of the *KAI2I* and *KAI2J* clades must have occurred relatively early in gymnosperm evolution, since both proteins are found in ginkgo and cycads, although *KAI2I* is not found in conifers (Figure 3; Figure 4). There are also many local duplications in the *KAI2* lineage, with some species having up to five *eu-KAI2* paralogs. However, the overall evolutionary trend in the *eu-KAI2* clade (as also suggested by the generally short branch lengths), is one of conservation, rather than innovation (Figure 1, Figure 3).

**Figure 3:**
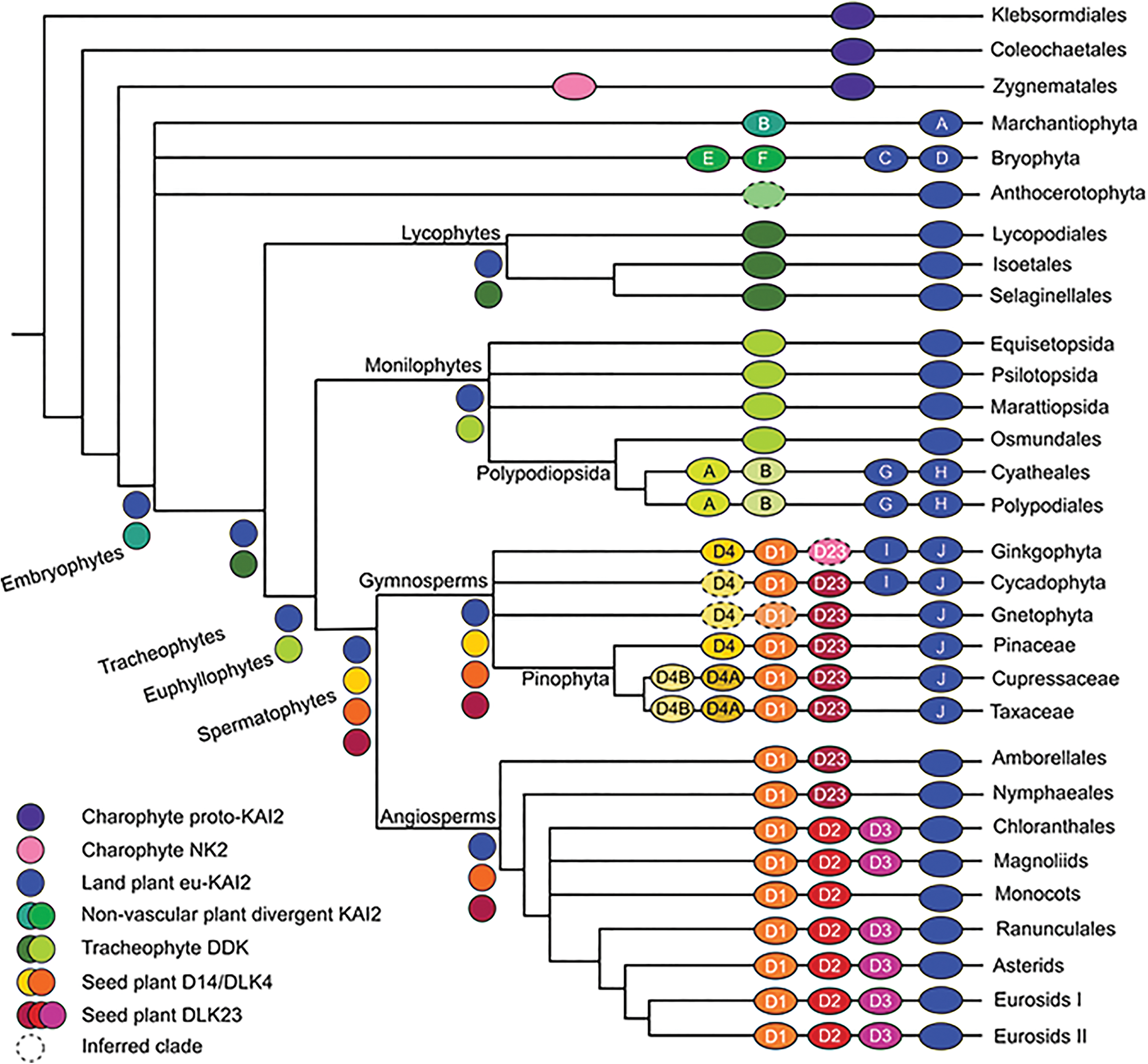
Reconstruction of D14/KAI2 family evolution. Schematic depicting the complement of D14/KAI2 proteins in major land groups, and their inferred evolutionary origin. Each branch indicates a major land plant group; lycophytes, monilophytes and gymnosperms are further sub-divided into relevant orders/families/etc. The ovals on each branch indicate the core complement of proteins in that group or sub-group, and are coloured according to the scheme indicated at the bottom left. Clades which are inferred by parsimony are denoted with a hatched line. Letters and numbers in the ovals denote the clade name as outlined in Table 2. Letters and numbers in the circles indicate clade names. D1 = D14, D2 = DLK2, D3 = DLK3, D4 = DLK4, D23 = DLK23. Circles without symbols at internal branching points represent the minimum inferred D14/KAI2 protein complement in the last common ancestor of each major land plant group.

**Figure 4:**
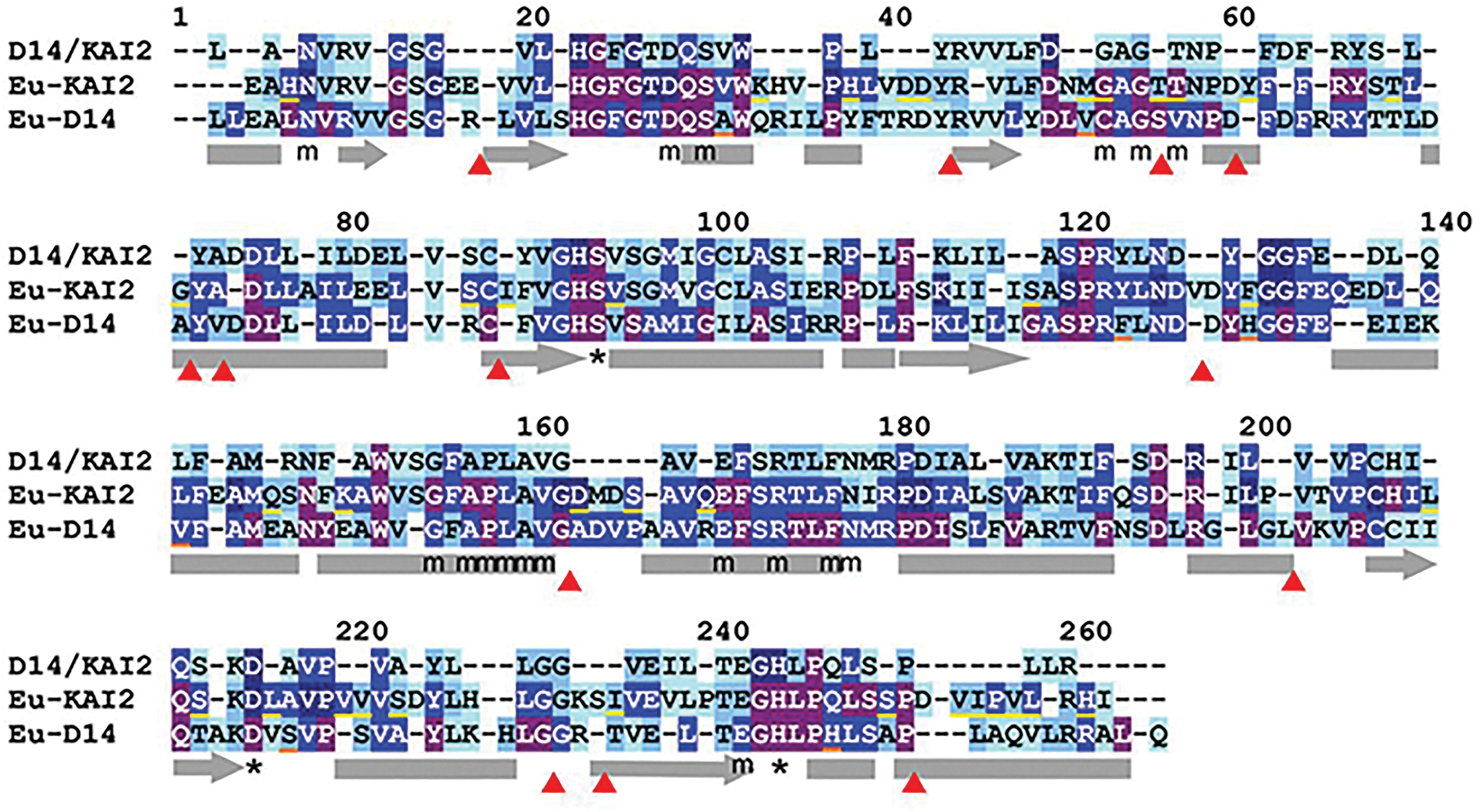
Eu-KAI2 proteins have highly conserved structure. Alignment illustrating conservation of primary protein structure in D14/KAI2 proteins. The 265 core positions (numbered) are shown in the alignment, for the whole family (top row), for eu-KAI2 proteins (middle row) and for eu-D14 proteins (bottom row). Positions where the same amino acid is present in >50% of sequences in the clade are denoted by corresponding letter; other positions are denoted by a dash. The colouring of each conserved residue indicates the degree of conservation; pale blue >50%, light blue >70%, mid-blue >90%, dark blue >99%, purple =100%. Structural features are annotated below the alignment. The catalytic triad is indicated by *. MAX2-interacting residues are indicated by a lower-case ‘m’. Predicted alpha helices (based on the crystal structure of AtKAI2 (PDB code 4HRX1A); are shown by grey bars, predicted beta strands by grey bars with an arrow. The discrete positions in the poly-peptide chain where insertions (or deletions) can be tolerated are illustrated with red arrow heads. Residues that are characteristic of eu-KAI2 proteins are underlined in yellow, residues characteristic of eu-D14 are underlined in orange (see Figure 5).

Conversely, the evolutionary history of the *DDK* clade is one of divergence and diversification. The liverwort and moss clades (*KAI2B*, *KAI2E*/*F*) are on relatively short branches (Figure 3) and have been categorized previously as encoding KAI2-like proteins. The lycophyte and monilophyte ‘DDK’ proteins are not obviously similar to the previously described KAI2, D14 or DLK2 protein types, nor indeed to each other. These clades also have long internal branch lengths, indicating a high degree of sequence divergence within the clades (Figure 3). In the leptosporangiate fern core group there has been a duplication in the *DDK* lineage, and the resulting DDKA and DDKB protein types are strongly divergent from both each other and from other monilophyte DDK proteins. In seed plants, there are a number of major duplications and evidence for significant innovation in protein sequence (Figure 3). In gymnosperms, we identified *eu-D14* sequences that form a sister clade to the well-characterized angiosperm *D14* clade. We also identified a second set of sequences in gymnosperms that are closely related to *D14*, which we named *DWARF14-LIKE4* (*DLK4*). These form a sister clade to the gymnosperm/angiosperm *eu-D14* clade, suggesting that the duplication that gave rise to *DLK4* occurred before the separation of gymnosperms and angiosperms (Figure 1). This in turn implies that the *DLK4* clade has been lost from angiosperms (Figure 3). Within the conifers there has been a major duplication in the *DLK4* lineage giving rise to two sub-clades (*DLK4A* and *DLK4B*); since *DLK4B* is not found in Pinaceae, the separation of *DLK4A* and *DLK4B* seems to post-date the divergence of pines and other conifers (Figure 3).

In angiosperms, we also discovered a third clade of proteins in addition to the expected *D14* and *DLK2* clades, which appeared as a sister clade to *DLK2* in our analysis; we named these sequences *DWARF14-LIKE3* (*DLK3*)(Figure 1). Although our phylogenetic reconstruction suggests that the separation of *DLK2* and *DLK3* occurred before the radiation of extant angiosperms, the distribution of *DLK3* sequences in our dataset suggests a slightly different history. We did not recover any *DLK3*-like sequences from the completed genome sequence of *Amborella trichopoda* (the sister group to all other angiosperms), nor from the plants in the other early-diverging angiosperm orders (Nymphaeales, Austrobaileyales). We did identify unambiguous *DLK3* sequences from the Chloroanthales and magnoliids, but not from any monocot species (including the fully sequenced genomes in Poaceae), despite extensive screening; we could however identify *DLK2* sequences from across the monocot group. *DLK3* sequences are present throughout the eudicots, though there have been sporadic losses, including in Brassicaceae. The exact inter-relationship of the major angiosperm lineages is currently uncertain, but one well-supported model is that monocots are sister to a clade containing magnoliids, Chloranthales and eudicots (Wickett et al, 2014). Under this scenario, the distribution of genes suggests that the separation of the *DLK2* and *DLK3* lineages occurred after the divergence of monocots and other angiosperms (Figure 3). Alternatively, *DLK3* could have been lost from the monocot lineage. We also identified a group of gymnosperm proteins that form a sister group to the combined angiosperm *DLK2*-*DLK3* clade, which we named *DLK23*. We also applied this name to the angiosperm proteins that pre-date the *DLK2*-*DLK3* split, and to the wider seed plant clade containing all these proteins (Figure 1, Figure 3).

### Sequence conservation among D14/KAI2 proteins

To further understand the consequences of the evolutionary trajectories of the *D14*/*KAI2* family members, we performed an in-depth analysis of their primary protein structure. Using our alignment, we identified a core set of 265 positions that occur in almost every D14/KAI2 protein (Figure 4). The start and end positions of the polypeptide chain vary between individual sequences, but the majority of sequences are within the range −15 to 280. Extra amino acids are inserted within the core of the protein in some sequences; these are usually located outside secondary structural elements such as α-helices (Figure 4). Most of these insertions are not conserved even between closely related sequences, although there are some exceptions. For instance, DDKB proteins from monilophytes have a conserved insertion of five amino acid after position 73.

In order to make comparisons across the family, we focused our attention on the core positions 1-265. We examined the amino acid frequency at each of these core positions, in different subsets of sequences, and used the data to understand patterns of conservation and divergence. We classify a position as ‘conserved’ if the same amino acid occurs in more than 50% of sequences in the subset, ‘well conserved’ if found in more than 70% of sequences, ‘highly conserved’ if found in more than 90% of sequences, and ‘invariant’ if found in more than 99% of sequences. Using this methodology on the D14/KAI2 family as a whole (339 sequences), we found that 68% of positions are conserved, with 18.5% being highly conserved (Figure 4). Of these, 17 positions (6.4%) are invariant, including the catalytic triad of serine, aspartate, and histidine (positions 94, 215, 244 respectively)(Figure 4, Table 3). Most of the highly-conserved residues cluster together in the polypeptide chain, forming motifs that are presumably important for protein activity (Figure 4).

### Eu-KAI2 clade members have strong sequence conservation

Using this approach, we tested the hypothesis that evolution in the *eu-KAI2* super-clade has generally been conservative. We analyzed amino acid frequencies from 127 eu-KAI2 proteins, and found that 22% of positions are invariant among eu-KAI2 proteins and 89% are conserved (Figure 4, Table 3). By comparison, in the DDK super-clade only 5.6% of positions are invariant, with 63% conserved (Table 3). Indeed, the level of conservation across eu-KAI2 proteins as whole is very comparable to conservation within taxon-level KAI2 clades. For instance, the angiosperm eu-KAI2 clade has 24% invariant positions and 94% conserved (Table 3). Together with the short branch lengths, the similarity in the level of between-clade and within-clade conservation in the eu-KAI2 super-clade supports the idea of a conservative evolutionary history.

Our dataset also allowed us to define a set of residues that are characteristic of eu-KAI2 proteins. We identified 39 positions where the same amino acid is present in at least 70% of eu-KAI2 sequences, and at which the same amino acid is present in less than 30% of DDK clade proteins (Figure 5A). These are not necessarily the best-conserved positions in eu-KAI2 proteins (Figure 4), but are those which are most characteristic of eu-KAI2 sequences. When compared to this reference set of residues, individual eu-KAI2 sequences from across the super-clade match at 35–38 out of 39 positions. Conversely, individual D14 sequences only match at 2–5 of these positions, for instance (Figure 5A, Supplementary Data 2). Eu-KAI2 proteins have therefore been generally well conserved through land plant evolution, which in turn implies conservation of eu-KAI2 function.

**Table 3:**
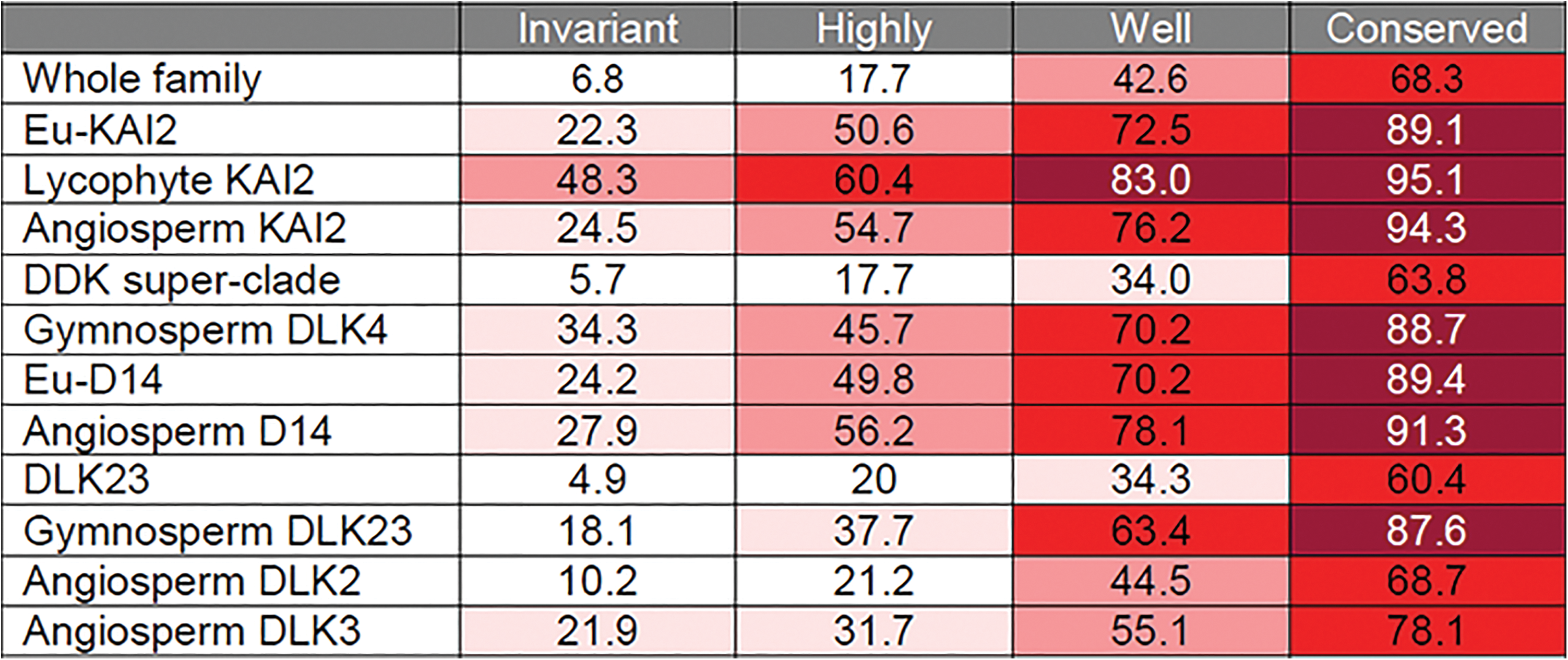
Protein sequence conservation in D14/KAI2 proteins. Table showing the degrees of protein sequence conservation in various D14/KAI2 clades. Four degrees of conservation were used: ‘invariant’ (>99% of sequences in a given clade have the same amino acid at a given position), ‘highly-conserved’ (>90% of sequences in a given clade have the same amino acid at a given position), ‘well-conserved’ (>70% of sequences in a given clade have the same amino acid at a given position) and conserved (>50% of sequences in a given clade have the same amino acid at a given position). The values in the table indicate the percentage of positions that fall into these categories in a given clade. So, for instance, 6.8% of positions are invariant in the whole family. Values are cumulative, so ‘conserved’ includes all the positions that are well-conserved, highly-conserved and invariant. Shading indicates values greater than 20% (light pink), 40% (dark pink), 60% (red) and 80% (dark red).

**Figure 5:**
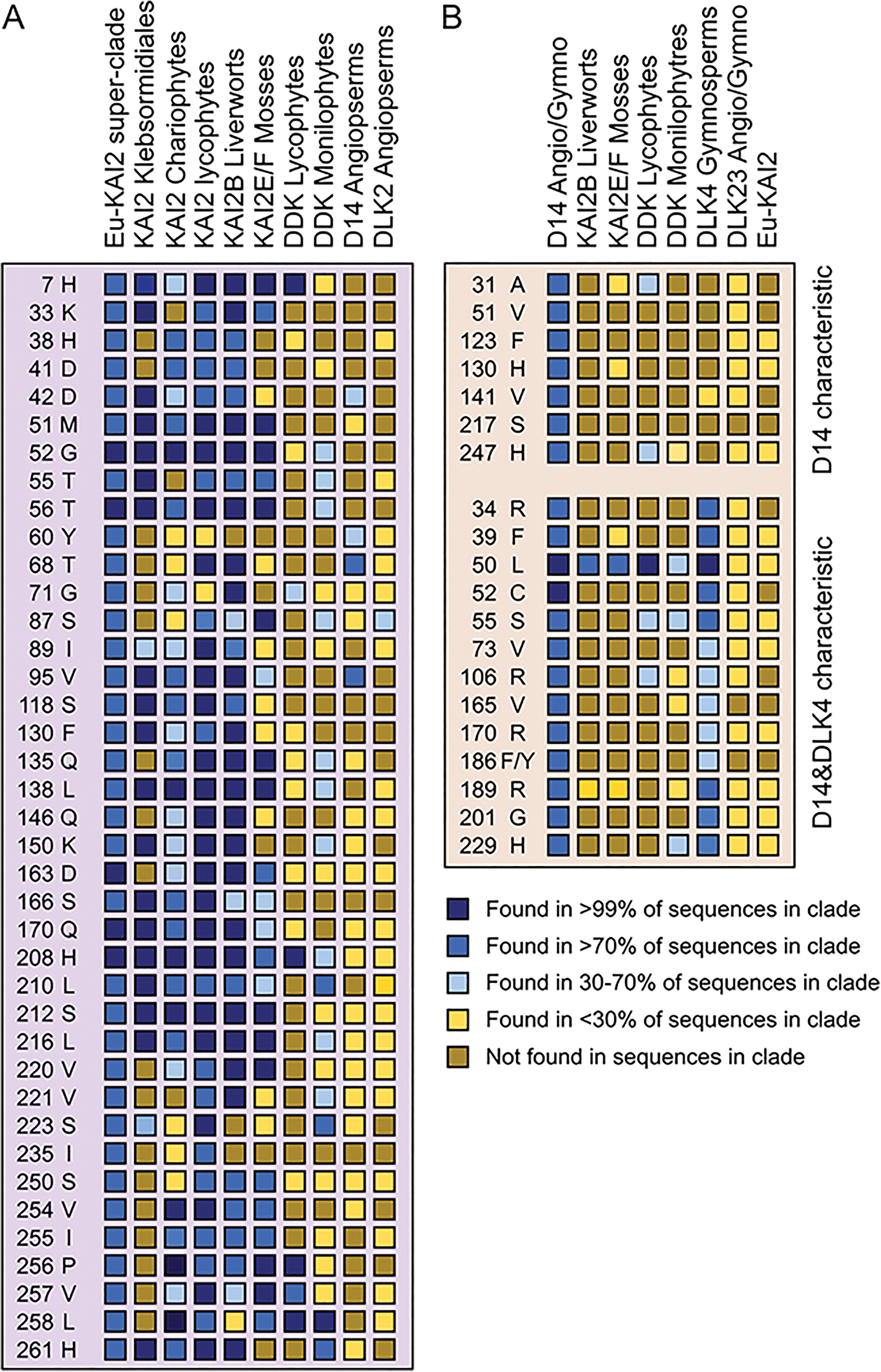
KAI2 and D14 protein characteristics. **A)** We identified well-conserved positions in eu-KAI2 proteins (i.e. >70% of sequences have the same amino acid) in which the amino acid is characteristic of eu-KAI2 proteins (i.e. found in <30% of other sequences). These are listed at the left (position and amino acid). We then tested whether various clades share elements of this structure (i.e. how frequently is the same amino acid is found at the same position in that clade). Charophyte and lycophyte KAI2 proteins are a close match, while KAI2B and KAI2E/F proteins from liverworts and mosses respectively have considerable similarity. However, DDK, D14 and DLK2 proteins do not share these characteristics. **B)** We performed the same analysis with eu-D14 proteins, but only identified 7 characteristic residues. We thus extended the search to the combined D14-DLK4 clade, and identified another 13 residues characteristic of the wider clade. These are listed at the left (position and amino acid). these elements. Very little conservation of these characteristic residues is found in other members of the DDK family.

### Charophyte D14/KAI2 family members may encode proto-KAI2 proteins

We examined the charophyte KAI2-like proteins relative to our eu-KAI2 reference set and found that they matched at 20–29 positions (Figure 5A; Supplementary Data 2). This suggests that while these proteins have relatively strong similarity with eu-KAI2 proteins, they are probably not true KAI2 proteins. To test this idea further, we generated homology models of charophyte KAI2 proteins using the crystal structure of karrikin-bound *Arabidopsis thaliana* KAI2 as a template (Guo et al, 2013). Focussing on the ligand binding pocket, we observed that some of the charophyte proteins have pockets similar to *A. thaliana* KAI2 (Figure 6A,I-L; Supplementary Data 3), while others had larger pockets. This difference is primarily determined by substitution of the ‘intrusive’ phenylalanine residue (F25) that limits the volume of the eu-KAI2 pocket for a leucine residue. These data are consistent with the idea that charophyte KAI2 proteins are similar to eu-KAI2 proteins, but do not completely conform to the conserved eu-KAI2 structure.

**Figure 6:**
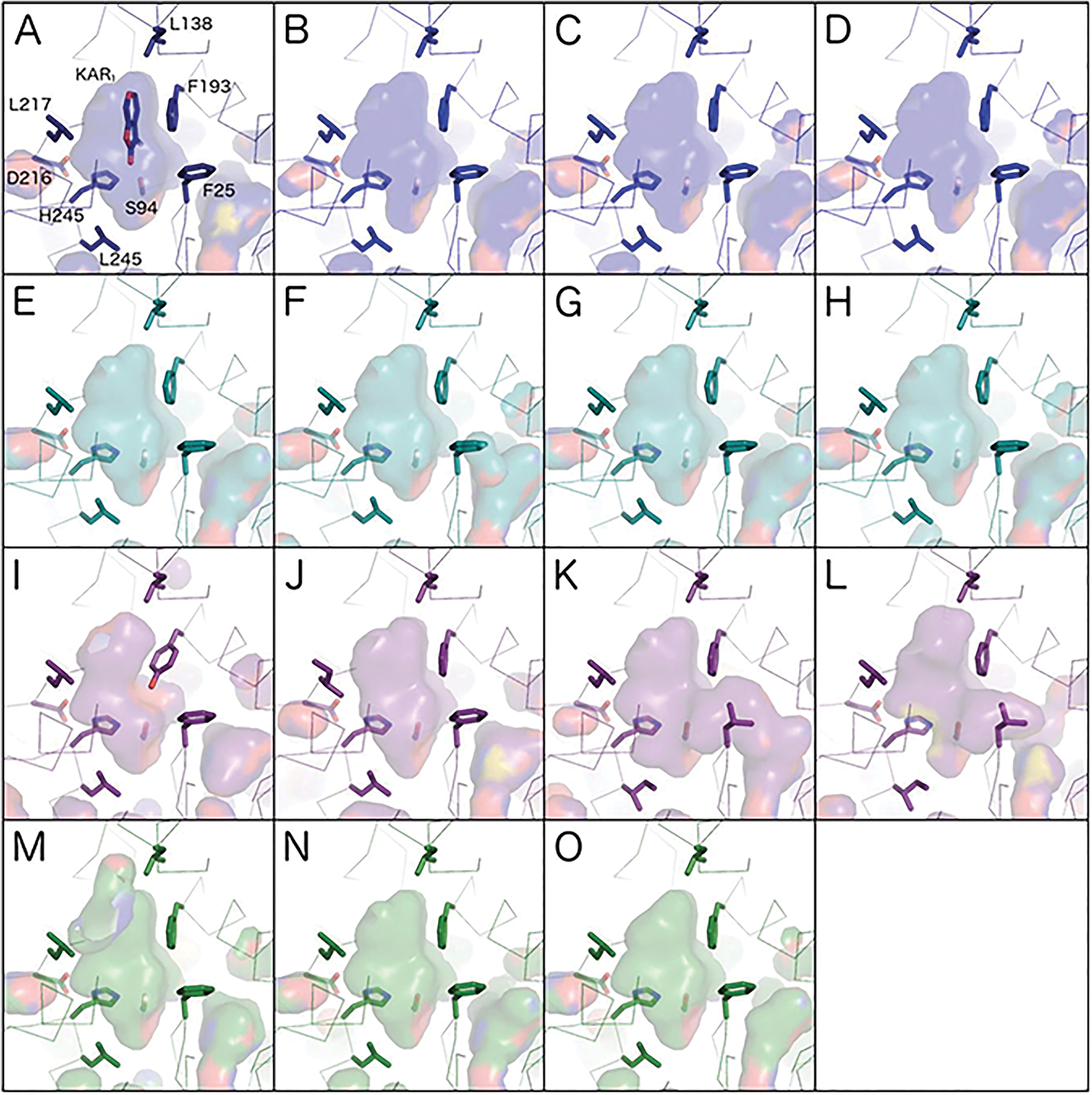
Homology models of KAI2 sequences. Models are shown in ribbon representation with the residues that influence the active site cavity shown in stick representation. Cavities are depicted as a transparent surface. Oxygen, nitrogen and sulphur atoms are coloured red, blue and yellow respectively. **(A)** The crystal structure of *Arabidopsis thaliana* KAI2 in complex with karrikin (KAR_1_) is shown in navy blue (PDB code 4JYM). Residue numbers correspond to the unified numbering scheme as in figure 5, which are −1 relative to those of *A. thaliana* KAI2. **(B-D)** Liverwort KAI2A homology models are shown in royal blue; **(B)** *Lejeuneaceae sp*. **(C)** *Lunularia cruciata*, **(D)** *Ptilidium pulcherrimum*. **(E-H)** Liverwort KAI2B models are shown in turquoise; (**E)** *Riccia berychiana*, **(F)** *Calypogeia fissa*, (**G)** *Lunularia cruciata*, (**H)** *Marchantia polymorpha*. **(I-L)** Charophyte KAI2 models are shown in purple; **(I)** *Klebsormidium subtile*, (**J)** *Chara vulgaris*, *Coleochaete scutata*, **(L)** *Coleochaete irregularis*. **(M-O)** Moss KAI2E/F models are shown in purple; **(M)** *Sphagnum recurvatum* KAI2E, (**N)** *Timmia austriaca* KAI2F, **(O)** *Tetraphis pellucida* KAI2F.

### Liverwort DDK clade members have conserved KAI2 structure

We next turned our attention to the DDK clade, which has lower overall amino acid conservation. We assessed whether the DDK proteins from liverworts (KAI2B), which have previously been characterized as KAI2-like, have conserved KAI2 features. We found that individual KAI2B proteins match the eu-KAI2 reference set at 29–33 out of 39 positions (Figure 5A; Supplementary Data 2). Although this is lower than eu-KAI2 proteins from liverworts (KAI2A), it suggests that these proteins could retain aspects of KAI2 primary protein structure. To test this idea, we generated homology models of liverwort KAI2B proteins using the crystal structure of karrikin-bound *Arabidopsis thaliana* KAI2 as a template (Guo et al, 2013). In each case, we found that the ligand binding pocket of KAI2B proteins are predicted to be essentially identical to Arabidopsis KAI2, and indeed liverwort KAI2A proteins (Figure 6A-H; Supplementary Data 3). Thus, while KAI2B proteins may be somewhat divergent relative to eu-KAI2 proteins, they probably still retain key features of eu-KAI2 structure.

### Moss and lycophyte DDK clade members do not have KAI2- or D14-like sequences

Conversely, when we analyzed the moss KAI2E and KAI2F proteins, we found that they only matched the KAI2 reference set at 22–24 positions (Figure 5A, Supplementary Data 2). This is a more considerable divergence from eu-KAI2 than liverwort KAI2B proteins, and could imply a corresponding alteration in function. Indeed, structural modelling of the KAI2E/KAI2F proteins from *P. patens* has previously suggested that some of these proteins have altered ligand-binding pockets relative to eu-KAI2 proteins in the same species (Lopez-Obando et al, 2016). However, modeling of newly available KAI2E/F sequences from other mosses did not suggest an major divergences from the KAI2 binding pocket (Figure 6M-O).

When we analyzed lycophyte DDK proteins, we found that they had much less affinity with eu-KAI2 proteins, matching the reference set at only 5–10 positions (Figure 5A; Supplementary Data 2). To test whether any of these proteins have signatures of D14-type SL receptors, we tried to identify a reference set of D14-characteristic amino acids comparable to our KAI2 reference set. We identified 13 positions at which the same amino acid is present in more than 70% of proteins in the DLK4/D14 clade, and is found at the same position in less than 30% of sequences in both the eu-KAI2 clade and in the wider DLK23 clade (since none of these proteins are currently considered to be an SL receptor)(Figure 5B). We also identified a further 7 positions with amino acids characteristic of eu-D14 proteins alone (Figure 5B). Known D14 proteins typically match this reference set at 15-20 out of 20 positions (Supplementary Data 2). When we compared individual KAI2E/F proteins to this reference set, they matched at only 0–4 positions (Figure 5B; Supplementary Data 2). Similarly, lycophyte DDK proteins only matched the D14 reference set at 1–3 positions. Neither of these types of protein thus displays particular similarity to known strigolactone receptors at the level of primary protein sequence. Furthermore, lycophyte DDK proteins display little specific similarity to any characterized member of the D14/KAI2 family. Consistent with this, homology models of lycophyte DDK proteins predicted ligand binding pockets that were neither KAI2-like nor D14-like (Figure 7A-D; Supplementary Data 3).

**Figure 7:**
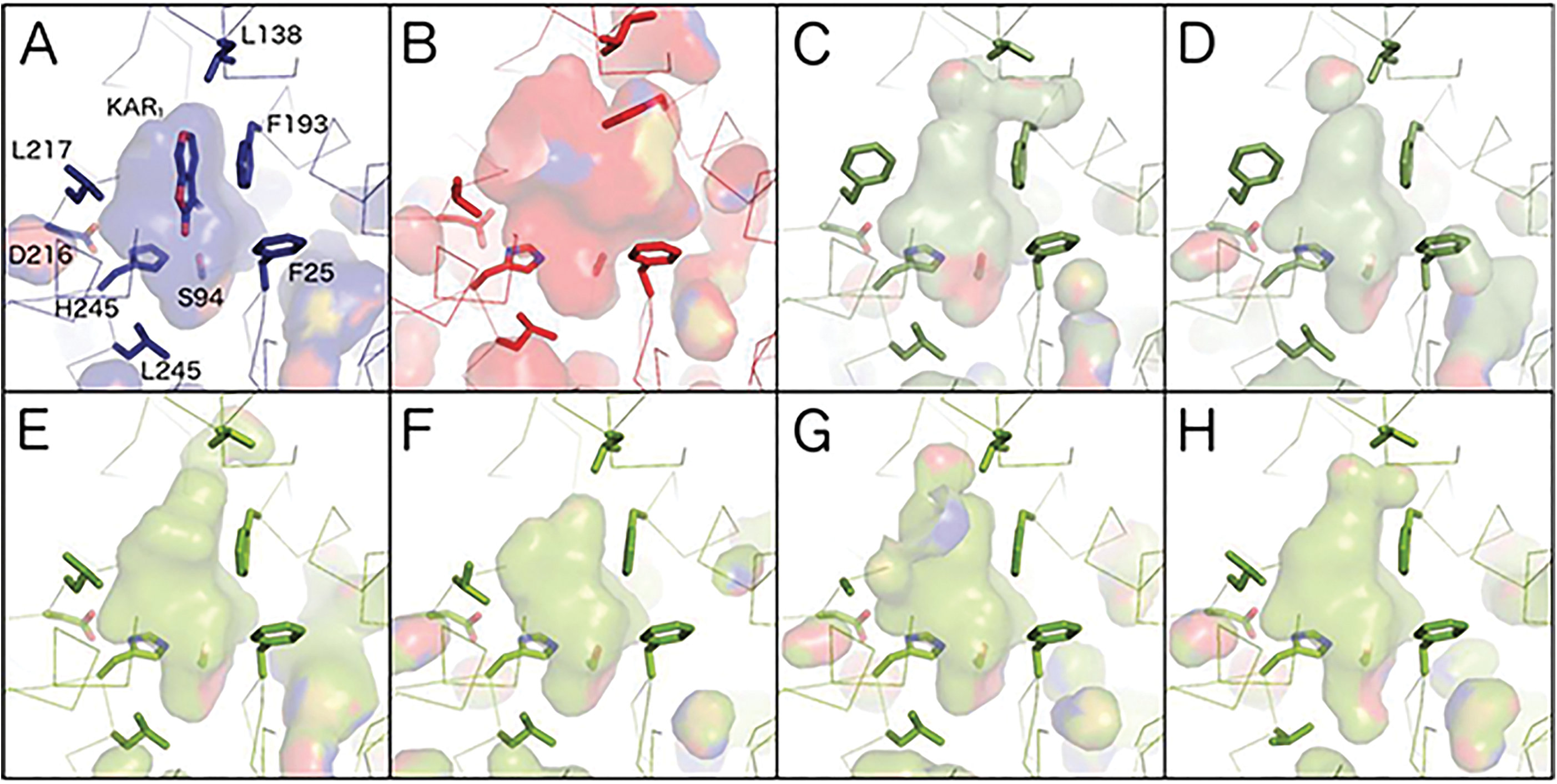
Homology models of DDK sequences. Models are shown in ribbon representation with the residues that influence the active site cavity shown in stick representation. Cavities are depicted as a transparent surface. Oxygen, nitrogen and sulphur atoms are coloured red, blue and yellow respectively. **(A)** The crystal structure of *A. thaliana* KAI2 in complex with karrikin (KAR_1_) is shown in navy blue (PDB code 4JYM). Residue numbers correspond to the unified numbering scheme as in figure 5, which are −1 relative to those of *A. thaliana* KAI2. **(B)** The crystal structure of *A. thaliana* D14 is shown in red. **(C-D)** Lycophyte DDK homology models are shown in olive green; **(C)** *Selaginella moellendorfii* (previously referred to as KAI2b) **(D)** *Lycopodium annotinum*. **(E-H)** Monilophyte DDK homology models are shown in lime green; **(E)** *Osmunda* sp. DDKb, **(F)** *Polypodium amorphum* DDKA, **(G)** *Cystopteris fragilis* DDKA, **(H)** *Asplenium platyneuron* DDKB.

### Seed plant DLK23 and monilophyte DDK proteins may function independently of MAX2

Recent work has delineated the residues in D14 that are needed for interaction with MAX2-class F-box proteins (Yao et al, 2016). We confirm that these 18 residues are strongly conserved in D14 proteins, as suggested by Yao et al (2016). We also noted that 16 of those residues are very highly conserved in the eu-KAI2 super-clade, strongly suggesting that KAI2 proteins interact with MAX2 proteins through exactly the same interface as D14 (Table 4). However, the level of conservation across the D14/KAI2 family as a whole is considerably lower than in either D14 or KAI2 groups, and we thus examined conservation of MAX2-interaction positions in other clades. Remarkably, we observed that of these 18 positions, 12 were not conserved in the highly divergent monilophyte DDKA and DDKB clades; 6 of these positions were not conserved in any monilophyte DDK protein (Table 4). Similarly, we found that 5, 7 and 8 of these positions are not conserved in DLK23, DLK2 and DLK3 proteins respectively (Table 4). Curiously, 5 of these positions are not conserved in the DLK4B, despite the MAX2-interface being otherwise conserved in the wider D14/DLK4 clade. These data suggest that these proteins may act independently of MAX2 signalling; this idea is discussed further below.

The monilophyte proteins occupy an intermediate position in the DDK clade, and we had therefore expected they would have protein sequences intermediate between the KAI2-like proteins in liverworts and eu-D14 proteins in seed plants. Individual monilophyte DDK proteins match the KAI2 reference set at 5–13 positions and the D14 reference set at 0–5 positions (Figure 5), suggesting that, like lycophyte DDK proteins, they are not especially similar to characterized proteins, and have unique structural features. Indeed, homology modelling suggests that these proteins have quite variable ligand binding pockets that are generally larger than eu-KAI2 proteins, but smaller than D14 proteins (Figure 7E-H, Supplementary Data 3). This is consistent with the general level of variation among monilophyte DDK proteins. Similarly, we observed that sequence conservation across the wider DLK23 clade is low; only 5% of positions are invariant, and only 60% conserved (Table 3). As would be expected, none of these proteins show affinity with KAI2 or D14 sequences (Figure 6A). It is therefore possible that loss of MAX2-interaction in these proteins has relaxed the structural requirements for protein function, resulting in divergent sequence characteristics.

**Table 4:**
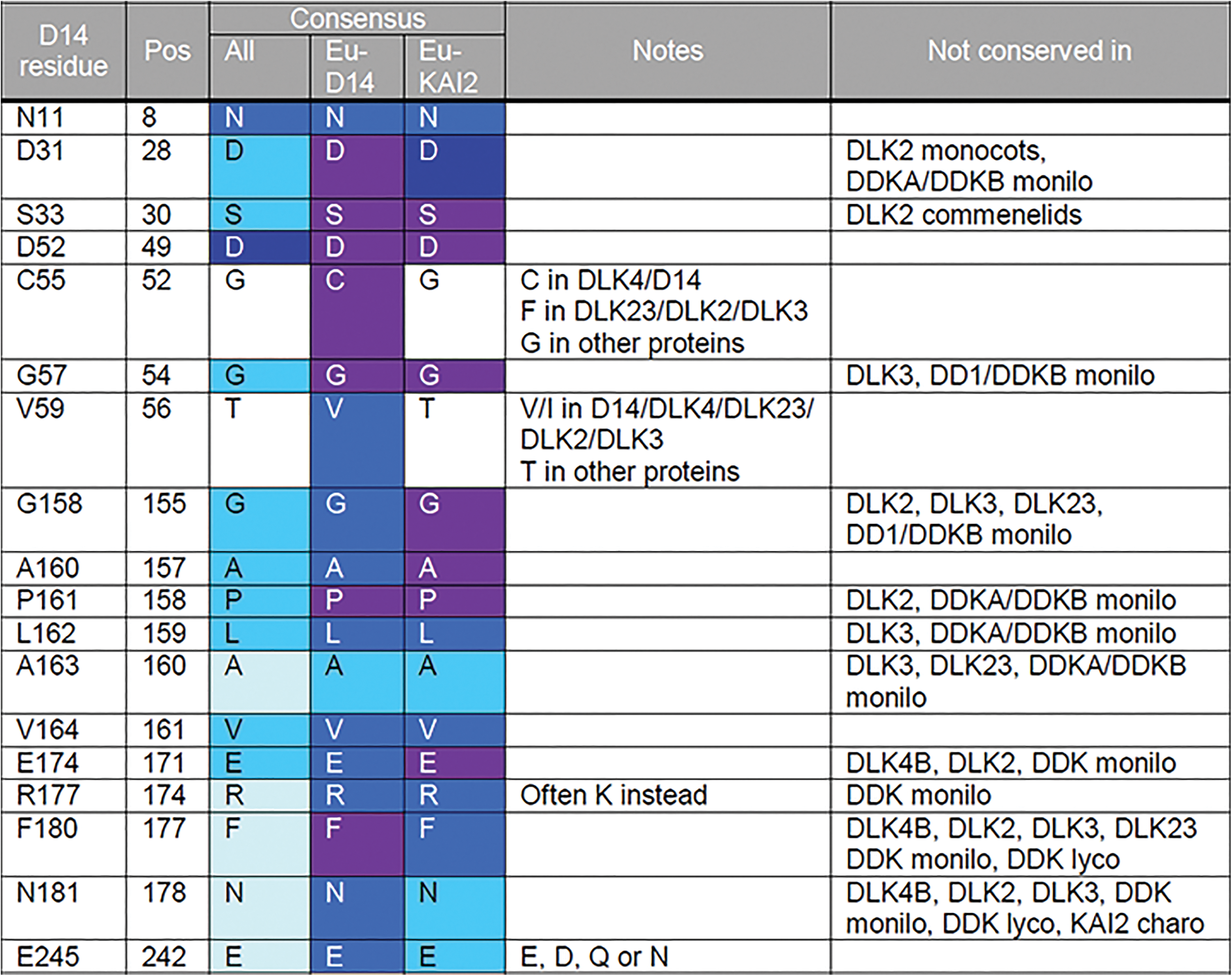
MAX2-interacting residues. The residues in the left-most column are those identified by Yao et al (2016) as playing a role in the interaction of Arabidopsis D14 with D3 (=MAX2) from rice. Numbers in the first column are relative positions with the AtD14 protein; these are corrected to our unified system in the second column. The consensus amino acids at those positions in the whole family, eu-D14 and eu-KAI2 clades are given in the next three columns. Shading indicates the degree of conservation at the position (pale blue >50%, light blue >70%, mid-blue >90%, dark blue >99%, purple 100%). The final column indicates clades in which these residues are not conserved.

## DISCUSSION

### KAI2 signalling is highly conserved

Previous studies showed that proteins resembling KAI2 are found throughout land plants and in charophyte algae (Delaux et al, 2012; Waters et al, 2012, Waters et al, 2015). Consistent with this, we demonstrate that one of the two major clades in the land plant *D14*/*KAI2* family contains only sequences that strongly resemble Arabidopsis KAI2. We demonstrate with very high resolution that these eu-KAI2 proteins are exceptionally conserved in protein sequence across the clade. Eu-KAI2 proteins have a clearly definable primary protein structure that is distinct from other members of the D14/KAI2 family, and their high levels of conservation arises from both shared-ancestral and shared-derived characteristics (Figure 5, Figure 6). These data strongly suggest that there are very specific structural requirements for KAI2 function, and that these functional characteristics have been conserved throughout land plant evolution. Our results demonstrate that D14/KAI2 family proteins from charophytes do not quite meet the definition of eu-KAI2 proteins, but that they do have significant similarity with KAI2 proteins; we have thus categorized them as proto-KAI2. While the function and role of D14 in SL signalling is well-understood, KAI2 proteins represent an enigma. In Arabidopsis, KAI2 is required for perception of karrikins, but has clearly defined developmental roles that are unrelated to karrikins; nor is Arabidopsis a naturally fire-following species (Waters et al, 2012; Bennett et al, 2016). This has led to the hypothesis that KAI2 regulates development in response to an unknown endogenous ligand (KL), which is mimicked by karrikins (Flematti et al, 2013; Conn and Nelson, 2015). Consistent with an ancestral role of KL perception, expression of the eu-KAI2 protein from *Selaginella moellendorffii* (SmKAI2A), can partially rescue an Arabidopsis *kai2* mutant, but does restore perception of karrikins (Waters et al, 2015). Identification of KL itself will be an important step in understanding the conserved function of KAI2 signalling across land plants (Sun et al, 2016).

### An early origin for strigolactone signalling?

Previous analyses of the *D14*/*KAI2* family have suggested that the origin of D14-type SL receptors is relatively recent, occurring within the vascular plant lineage, and perhaps restricted to seed plants (Delaux et al, 2012; Waters et al, 2012; Waters et al, 2015). Since SL sensitivity seems to be a widespread phenomenon in land plants and perhaps charophytes, this has led to significant speculation that non-canonical SL perception mechanisms exist in non-vascular plants (Bennett & Leyser, 2014; Waldie et al, 2014). For instance, it has been suggested that KAI2 proteins could act as SL receptors in mosses and liverworts (Bennett & Leyser, 2014). Our analyses show that, as far as a distinct primary protein structure can be defined for eu-D14, such proteins do indeed only exist in seed plants. However, the separation of the *DDK* clade (of which eu-D14 proteins are members) from the *eu-KAI2* clade occurred much earlier than previously suspected, at the base of the land plants. This raises the possibility that SL receptors might be a much earlier innovation in the D14/KAI2 famuily than previously suspected. The DDK protein from *Selaginella moellendorffii* (previously referred to as KAI2b) can hydrolyze SL-like stereoisomers of *rac*-GR24 (Waters et al, 2015), suggesting that it acts as a SL receptor. We show here that DDK proteins from lycophytes have little specific similarity to D14, which in turn suggests that other proteins in the clade could act as SL receptors despite their non-D14-like structure. However, understanding exactly when SL perception arose in the DDK lineage is contingent on understanding the evolution of land plants themselves. Although the phylogeny of vascular plants is well-established, there is still considerable debate regarding the relationship of non-vascular plants both to each other, and to vascular plants. Depending on which scenario is correct, our understanding of the evolution of SL signalling may be considerably altered.

The ‘traditional’ land plant phylogeny suggests that liverworts, mosses and hornworts form a grade with regard to vascular plants (e.g. Qiu et al, 2006). If this is correct, then the divergence of the eu-*KAI2* and *DDK* lineages would have occurred at the very base of the land plant tree (Figure 8A). Although slightly divergent in their general structure, liverwort KAI2B proteins appear to have the same ligand binding pockets as eu-KAI2 proteins (Figure 6). This is consistent with data showing that the KAI2B protein from *Marchantia polymorpha* preferentially hydrolyses non-natural stereoisomers of *rac*-GR24, rather than the SL-like stereoisomers (Waters et al, 2015). Indeed, it is currently unclear whether liverworts synthesize or perceive SLs (Waters et al, 2017). Under this model of land plant evolution, the evolution of SL perception could be envisaged to have occurred by gradual neo-functionalization of the *DDK* lineage (Figure 8A). Consistent with this, while KAI2B proteins are structurally similar to eu-KAI2 proteins, the moss proteins in the DDK lineage (KAI2E/F) are more divergent. There is clear evidence for SL perception in *P. patens*, and in this context, it is very interesting to note that a sub-set of *P. patens* D14/KAI2 proteins have previously been predicted to have SL-like ligand binding pockets (Lopez-Obando et al, 2016). All those proteins (KAI2Ea, KAI2Eb, KAI2Fd, KAI2Fe) are members of the *DDK* super-clade in our analysis. However, not all KAI2E/KAI2F proteins from *P. patens* are predicted to have divergent binding pockets (Lopez-Obando et al, 2016), and KAI2-like binding pockets were predicted in KAI2E/F proteins from other mosses (Figure 6). The status of KAI2E/KAI2F proteins as SL receptors is thus far from certain, and more work is needed to firmly establish their structure and function.

**Figure 8:**
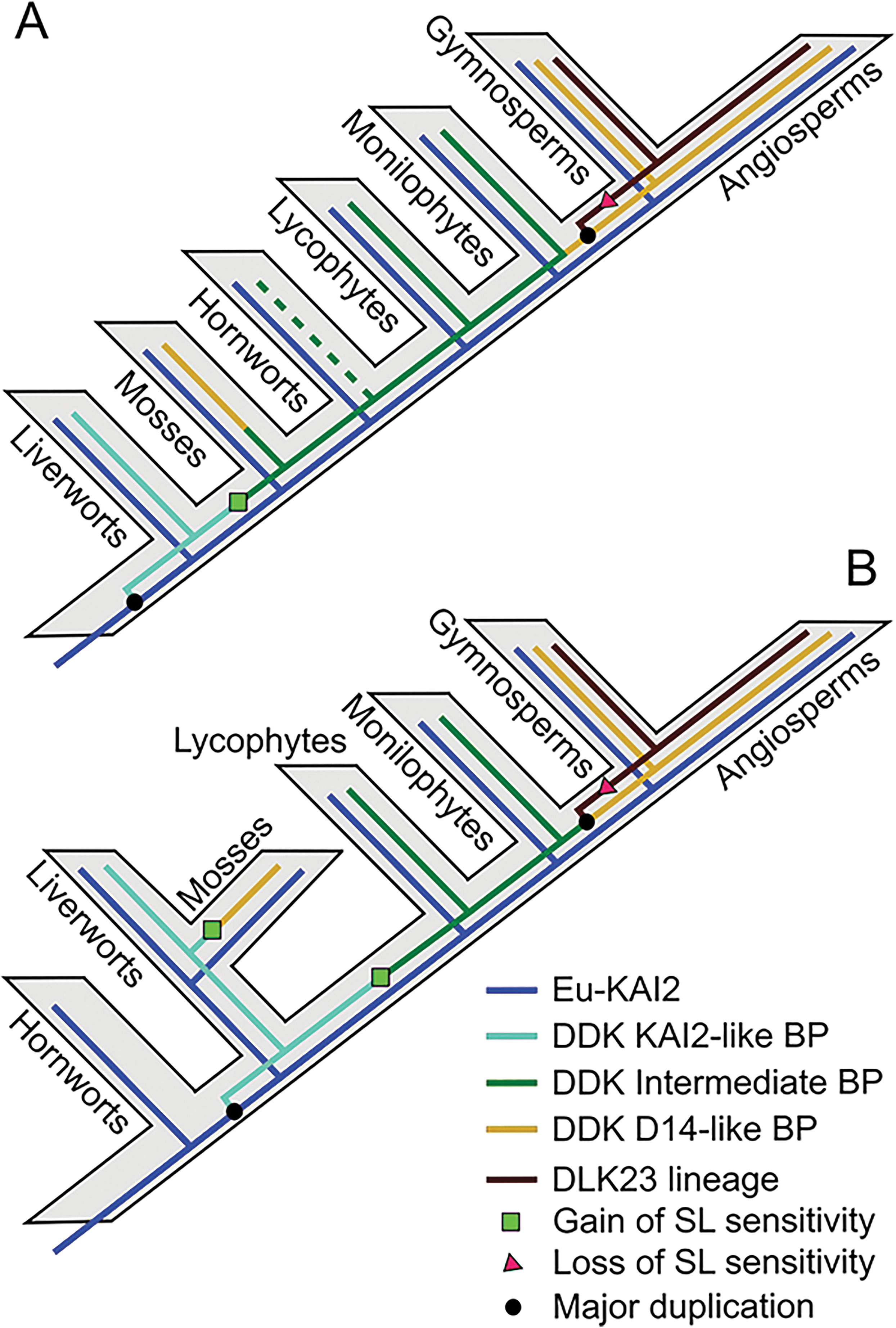
Models of *D14/KAI2* evolution. BP = binding pocket A) Traditional model of land plant evolution, with evolution of the *D14*/*KAI2* family superimposed. A single origin of SL perception in the *DDK* lineage would be sufficient to explain known patterns of SL sensitivity. B) ‘Hornworts-basal’ model of land plant evolution, with evolution of the *D14*/*KAI2* family superimposed. Two independent origins of SL perception in the *DDK* lineage would be required to explain known patterns of SL sensitivity.

A more recent model of land plant evolution suggests that hornworts are the earliest-diverging group of land plants, and that liverworts and mosses form a clade that is sister to vascular plants (Figure 8B)(Wickett et al, 2014). The ‘hornworts-basal’ model is controversial, but consistent with it, we only identified a single clade of KAI2-like proteins from hornworts, which in some of our analyses place this clade as a sister-clade to all other land plant *D14*/*KAI2* sequences (Figure 1, Figure 2) This would suggest that the duplication that created the *eu-KAI2* and *DDK* lineages occurred after the separation of hornworts from all other land plants (Figure 8B), although it should be noted that the recovery of a single hornwort clade could be due to the limitations of transcriptome databases. The close relationship of liverworts and mosses in this model (irrespective of their placement relative to hornworts) also has major implications for understanding the evolution of SL signalling. If this scenario is correct, then liverwort KAI2B and moss KAI2E/F are probable sister clades. Given the eu-KAI2 like structure of KAI2B protein, this would firmly imply that the ancestral state in the joint KAI2B-E/F clade would involve a KAI2-like binding pocket. If moss KAI2E/F proteins do indeed act as SL receptors, this would mean that SL-like binding pockets would have evolved twice independently in the *DDK* lineage, in mosses and vascular plants (Figure 8B).

Our ability to precisely understand the origins of SL perception in the *DDK* lineage is thus currently limited by the lack of clarity regarding non-vascular plant phylogeny. It is nevertheless clear that the evolutionary trajectory of the *DDK* lineage is away from an initially KAI2-like structure, and that SL perception probably arose in the lineage at the latest in vascular plants. Given the high conservation between eu-KAI2 proteins, it is therefore very likely that that the majority of proteins in the DDK lineage are at least neo-functional with respect to KAI2. The primary question is thus whether they are neo-functional as SL receptors, or as something rather different. Our data suggest that the structural requirements for SL perception in vascular plants may be relatively relaxed, and even eu-D14 proteins only have limited shared-derived characteristics (Figure 5A). We speculate that interactions with protein partners (such as SMXL proteins) may have driven the evolution of D14-like structure, rather than requirements for SL perception in itself.

### MAX2-coupled signalling in the D14/KAI2 family

Alongside the origin of specific SL receptors, the evolution of SCF^MAX2^-coupling with D14/KAI2-signalling has also been a subject of debate. Two points have been emphasized; first, that proto-KAI2 proteins are present in charophyte algae, but that MAX2 homologues do not seem to be (Challis et al, 2013; Delaux et al, 2012). Second, *P. patens max2* mutants are reported to have a very different phenotype relative to *P. patens* SL synthesis mutants (no filamentous growth versus excessive filamentous growth), suggesting that they are not in the same pathway (Proust et al, 2011; de Saint Germain et al, 2013). On this basis, it has been suggested that SL signalling in non-vascular land plants might proceed by non-canonical mechanisms (Bennett & Leyser, 2014; Waters et al, 2017). Our data provide us with some insights in this respect. Firstly, the defined MAX2-interaction interface found in D14 is highly conserved across most of the D14/KAI2 family, including in both eu-KAI2 and DDK proteins from liverworts, mosses, and hornworts. It therefore seems likely that these proteins do indeed signal via MAX2 in non-vascular plants. We thus hypothesize that the reported *max2* phenotype in *P. patens* arises from a lack of eu-KAI2 signalling, which in turn prevents expression of the SL-deficiency phenotype that would otherwise occur. Furthermore, our data show that the MAX2-interface is also conserved in charophyte D14/KAI2 proteins, tentatively suggesting the existence of MAX2-coupled signalling outside land plants. Thus far, a complete genome sequence is available for only one charophyte, *Klebsormidium flaccidum*. On this basis, it is not currently possible to conclude that MAX2 sequences are absent from all charophytes, rather than only absent from some charophytes.

In contrast to the strong conservation of D14 and KAI2 proteins, we identified several clades of proteins (DLK2 and DLK3 from angiosperms, DLK23 and DLK4B from gymnosperms, DDKA/DDKB and probably all DDK proteins from monilophytes) that are strongly divergent at the positions that comprise the MAX2-interface. The strong conservation between KAI2 and D14 proteins, which are both known to signal through MAX2, strongly implies that the amino acid composition of the MAX2-interface is critical. Furthermore, the strong conservation of these positions within the eu-KAI2 clade strongly implies that the cognate interaction surface on MAX2 has not significantly altered throughout the evolution of the land plants. On this basis, it seems a likely conclusion that the divergent proteins listed above do not interact with MAX2 (or that if they do, they do so in a radically different way to D14/KAI2). The possible counter-argument that these proteins might interact with specialized versions of MAX2 is demonstrably not the case in angiosperms, which usually have a single MAX2 paralog (Challis et al, 2013).

### A diversity of small molecular receptors?

The DLK23/DLK2/DLK3 clade remains the most enigmatic set of proteins in the D14/KAI2 family. Not only do they probably lack the conserved MAX2 interface, but they also have no known function, and are highly divergent from other D14/KAI2 proteins. DLK2 in Arabidopsis does not seem to be a receptor for SL or KL, at least as far as can be defined genetically (Waters et al, 2012; Bennett et al, 2016). One possibility is that the DLK23, DLK2, and DLK3 proteins act as receptors for a novel ligand, or perhaps multiple ligands. The wider DLK23 lineage in angiosperms has long internal branches, coupled with a lack of sequence conservation, but there is little evidence of gene loss. This suggests that the high degree of divergence does not simply represent drift in obsolete sequences. Rather, it may indicate continued innovation in the function of DLK23 proteins throughout angiosperm evolution, including the sub- or neo-functionalization process that led to independent DLK2 and DLK3 lineages. Since the pre-duplication DLK23 proteins from angiosperms tend to group with eu-DLK2 species in phylogenetic analyses, this tentatively suggests that DLK2 maintained the original structure/function of DLK23, and that the DLK3 lineage is neo-functionalized.

In addition to the DLK23 lineage, the fast-evolving DDK super-clade might contain further receptors for non-SL/KL ligands. For instance, since gymnosperms maintain conserved D14-type receptors, it is plausible that DLK4 proteins (and especially the more divergent DLK4B proteins) are not SL receptors. The same may be true of the highly divergent DDKA/DDKB proteins in monilophytes. Our work helps to broaden the structural biology platform for D14/KAI2 family members, and future work should provide very interesting insights into ligand-binding, structure and function if these diverse proteins, as well as their interactions with other SL signalling components.

## MATERIALS & METHODS

### Bioinformatic retrieval of D14/KAI2 sequences

Members of the *D14/KAI2* family were identified by BLAST searches against complete, annotated genomes from two major sources: Phytozome (www.phytozome.net), or the genome portals for individual species, for instance the Amborella genome project (www.amborella.org). BLAST searches were performed using the coding sequences of *Arabidopsis thaliana D14*, *KAI2* and *DLK2*, using the BLASTN option. Preliminary trees were assembled and used to guide the iterative interrogation of transcriptome databases, particularly those generated by the 1KP project (https://www.bioinfodata.org/Blast4OneKP/home). All sequences are listed in Supplementary Data 4. For transcriptome datasets, we BLASTed each major taxonomic group separately. Where novel protein types were identified within a taxon (e.g. Angiosperm *DLK3*) we re-BLASTed the same taxonomic group with the novel sequence, to increase the specificity of our searches. For non-annotated sequences from transcriptome datasets, we searched translations across all 6 reading frames to identify ORFs, and the longest ORFs were extracted for alignment.

### Alignment

Alignments were initially performed in Bioedit (Hall, 1997) using ClustalW (Thompson et al, 1994); *D14*/*KAI2* sequences are highly alignable, and this was a relatively trivial step. Full length sequences from completed genomes were used for the initial alignment, which was manually refined as necessary. We then added sequences from transcriptome databases, many of which are incomplete, but the alignment of full length sequences provided a scaffold to align these sequences correctly. The resultant alignment of 339 sequences is provided in Supplementary Data 5. For primary protein structure analyses, we focused on positions in the alignment that are present in most sequences. We removed the non-conserved extensions at the N- and C-terminus, producing an alignment with 265 core positions. We noted the positions of any non-conserved insertions within this core structure (Figure 4), and then removed them prior to the final analyses. This 795 nucleotide alignment/265 amino acid alignment was used for analyses of primary protein structure (Figure 4, Figure 5, Table 3, Table 4).

### Phylogenetic analysis

We performed preliminary phylogenetic analyses to explore the topology of the tree and the effect of inclusion or exclusion of various groups of sequences. We removed 15 nucleotides (5 positions; 57-60, 252) from the 795 nucleotide alignment that were not well conserved across all sequences, leaving a ‘maximum’ phylogenetic alignment of 780 nucleotides. We implemented nucleotide-level maximum likelihood analyses in PhyML (Guindon et al, 2010), and GARLI (Genetic Algorithm for Rapid Likelihood Inference; version 2.0) (Zwickl 2006), using the GTR+G+I model of evolution. These analyses are generally congruent with subsequent analyses, but identified some problems with tree reconstruction, particularly with respect to the position of charophyte and lycophyte KAI2 sequences.

For final analyses, the alignment was manually modified in AliView v1.18-beta7 (Larsson 2014), and areas of ambiguous alignment excluded from subsequent analyses. To determine the optimal model/partitioning scheme, we performed an exhaustive search in PartitionFinder v1.1.1 (Lanfear et al. 2012), with each of the three codon positions permitted its own parameters. All models were assessed, branch lengths were constrained to be proportional across partitions, and the topology was fixed to that inferred by a preliminary GARLI v2.01 (Zwickl 2006) analysis with each codon position given its own GTR+I+G model and rates permitted to vary across partitions; the optimal scheme was selected by the AIC (Akaike 1974). Maximum likelihood tree searches were performed under this model (codon positions 1 and 2 with their own GTR+I+G sub-models and codon position 3 with a TVM+I+G submodel; average rates permitted to vary across partitions) using GARLI v2.01 (Zwickl 2006), in the CIPRES Science Gateway (Miller et al. 2010). The GARLI tree searches were performed under the default settings with the exception that genthreshfortopoterm was increased to 40000; these searches were performed from 48 different random addition sequence starting trees. Support was assessed with 528 bootstrap replicates in GARLI, under the same settings as the best-tree searches, but with each bootstrap search performed from 24 different random addition sequence starting trees. The resulting bootstrap support values were mapped onto our ML phylogeny using the SumTrees v3.3.1 program in the DendroPy v3.12.0 package (Sukumaran and Holder 2010).

### Assessing tree robustness

We performed multiple analyses to test the robustness of our phylogenetic reconstructions, particularly the placement of KAI2B from liverworts and KAI2E/F from mosses within the DDK clade. Firstly, we removed each DDK clade from the alignment in turn, and re-ran the phylogenetic analysis in PhyML (Supplementary Data 1D). The ten recovered trees have four commonalities: 1) KAI2B is always placed in the eu-KAI2 lineage (except in the ‘No KAI2’ tree), 2) the rest of the DDK clade is always stably grouped together (although there are some variations in the exact branching order within the clade), 3) the relative position of KAI2E/F is completely invariant (except in the ‘No KAI2E/F’ tree, and 4) all of the trees place the eu-KAI2 lineage as a grade leading to the DDK clade. This latter point demonstrates that none of these trees are plausible in themselves, since the angiosperm eu-KAI2 clade is placed as a sister clade to the DDK clade containing moss, lycophyte, monilophyte, gymnosperms and angiosperm sequences. Secondly, we ran the analysis on an alignment cut down to match that of Waters et al (2012), using additional RbsQ (bacterial sigma factors with similarity to D14/KAI2 proteins) sequences identified in that study. If we rooted the resulting tree with RbsQ sequences, we observed the same basic topology as Waters et al (2012). However, if we rooted with *Selaginella moellendorfii* KAI2, we obtained the same basic topology as our main analyses, albeit with RbsQ as an in-group in the DDK lineage. Our analysis is thus congruent with the previous analysis of Waters et al (2012).

### Protein homology modelling

KAI2 and DDK sequences were modelled using the SWISSMODEL server (http://swissmodel.expasy.org) based on a multiple sequence alignment of KAI2 and DDK sequences (Bordoli et al. 2009). Numerous KAI2 crystal structures were available for use as a model template (Kagiyama et al. 2013, Zhao et al. 2013, Guo et al. 2013, Bythell-Douglas et al. 2013), however we chose the karrikin-bound *A. thaliana* structure (PDB code 4JYM)(Guo et al 2013) as it was the most informative for probing the regions of the protein involved in ligand interaction. Modelled sequences share 37-71% sequence identity with *A. thaliana* KAI2 as computed by BioEdit (Hall, 1997)(Supplementary Data 3). Protein structure and homology model figures were generated with PYMOL (DeLano 2002). Cavities within homology models were visualised using surface mode on the setting “Cavities & Pockets (Culled)” within PYMOL. Volume calculations were performed using the CASTp protein server (Dundas et al. 2006) using a probe radius of 1.4 Å. Initial calculations of volume misleadingly included regions of the surface of the protein adjacent to the cavity opening. This problem was circumvented by artificially blocking the cavity opening with a free alanine residue, which was not covalently attached to the protein molecule. This alanine was placed in the same *xyz* coordinates for all superposed homology models and crystal structures.

#### ACKNOWLEDGMENTS

This work was supported by grants from the European Research Council (N° 294514 – EnCoDe) and from the Monell Foundation to Dennis Stevenson. We gratefully acknowledge the use of sequence data generated by members of the 1000 Plants (1KP) initiative, and in particular Michael Melkonian, Barbara Surek, Jim Leebens-Mack, Michael Deyholos, Douglas Soltis, Pamela Soltis, Anders Larsson, Lisa Pokory and Lisa DeGironimo.

## REFERENCES

Akaike H. 1974. A new look at the statistical model identification. IEEE Transactions on Automatic Control 19: 716–723

Al-Babili S, Bouwmeester HJ. 2015. Strigolactones, a novel carotenoid-derived plant hormone. Annu Rev Plant Biol. 66:161–186.

Bennett, T., and Leyser, O. (2014). Strigolactone signalling: standing on the shoulders of DWARFs. Curr. Opin. Plant Biol. 22:7–13.

Bennett T, Brockington SF, Rothfels C, Graham SW, Stevenson D, Kutchan T, Rolf M, Thomas P, Wong GK, Leyser O, et al. 2014. Paralogous radiations of PIN proteins with multiple origins of noncanonical PIN structure. Mol Biol Evol. 31:2042–2060.

Bennett T, Liang Y, Seale M, Ward S, Müller D, Leyser O. 2016. Strigolactone regulates shoot development through a core signalling pathway. Biol Open. pii: bio.021402

Bordoli L, Kiefer F, Arnold K, Benkert P, Battey J, Schwede T. 2009. Protein structure homology modeling using SWISS-MODEL workspace. Nat Protoc 4:1–13.

Borghi L, Liu GW, Emonet A, Kretzschmar T, Martinoia E. 2016. The importance of strigolactone transport regulation for symbiotic signalling and shoot branching. Planta. 243:1351–1360

Bythell-Douglas R., Waters MT, Scaffidi A, Flematti GR, Smith SM, Bond CS. 2013. The structure of the karrikin-insensitive protein (KAI2) in Arabidopsis thaliana. PLoS ONE 8:e54758

Challis, R.J., Hepworth, J., Mouchel, C., Waites, R., Leyser, O. 2013. A role for more axillary growth1 (MAX1) in evolutionary diversity in strigolactone signalling upstream of MAX2. Plant Physiol 161:1885–1902.

Conn CE, Bythell-Douglas R, Neumann D, Yoshida S, Whittington B, Westwood JH, Shirasu K, Bond CS, Dyer KA, Nelson DC. 2015. Convergent evolution of strigolactone perception enabled host detection in parasitic plants. Science. 349:540–543.

Conn CE, Nelson DC. 2016. Evidence that KARRIKIN-INSENSITIVE2 (KAI2) Receptors may Perceive an Unknown Signal that is not Karrikin or Strigolactone. Front Plant Sci. 6:1219

Cox CJ, Li B, Foster PG, Embley TM, Civán P. 2014. Conflicting phylogenies for early land plants are caused by composition biases among synonymous substitutions. Syst Biol 63:272–279

DeLano W. 2002. The Pymol molecular graphics system. Schrödinger, LLC, San Carlos

Delaux PM, Xie X, Timme RE, Puech-Pages V, Dunand C, Lecompte E, Delwiche CF, Yoneyama K, Bécard G, Séjalon-Delmas N. 2012. Origin of strigolactones in the green lineage. New Phytol. 195:857–871

de Saint Germain A, Bonhomme S, Boyer FD, Rameau C. 2013. Novel insights into strigolactone distribution and signalling. Curr Opin Plant Biol. 16:583–589.

de Saint Germain A, Clavé G, Badet-Denisot MA, Pillot JP, Cornu D, Le Caer JP, Burger M, Pelissier F, Retailleau P, Turnbull C, et al. 2016. An histidine covalent receptor and butenolide complex mediates strigolactone perception. Nat Chem Biol. 12:787–794.

Doyle JA. Phylogenetic Analyses and Morphological Innovations in Land Plants. In: Ambrose BA, Purugganan M, editors. The Evolution of Plant Form. Blackwell. p. 1–50.

Dundas J, Ouyang Z, Tseng J, Binkowski A, Turpaz Y, Liang J. 2006. CASTp: computed atlas of surface topography of proteins with structural and topographical mapping of functionally annotated residues. Nucleic Acids Res. 34:W116–W118.

Flematti GR, Waters MT, Scaffidi A, Merritt DJ, Ghisalberti EL, Dixon KW, Smith SM. 2013. Karrikin and cyanohydrin smoke signals provide clues to new endogenous plant signalling compounds. Mol Plant. 6:29–37.

Flores-Sandoval E, Eklund DM, Bowman JL. 2015. A Simple Auxin Transcriptional Response System Regulates Multiple Morphogenetic Processes in the Liverwort Marchantia polymorpha. PLoS Genet. 11:e1005207.

Guindon S, Dufayard JF, Lefort V, Anisimova M, Hordijk W, Gascuel O. 2010. New Algorithms and Methods to Estimate Maximum-Likelihood Phylogenies: Assessing the Performance of PhyML 3.0. Systematic Biol. 59:307–321.

Guo Y., Zheng Z., La Clair JJ, Chory J, Noel JP. 2013. Smoke-derived karrikin perception by the α-hydrolase KAI2 from Arabidopsis. Proc. Natl. Acad. Sci. U.S.A. 110:8284–8289.

Hall T. 1997. BioEdit: Biological sequence alignment editor. Ibis Biosciences, Carlsbad

Hamiaux C, Drummond RS, Janssen BJ, Ledger SE, Cooney JM, Newcomb RD, Snowden KC. 2012. DAD2 Is an alpha/beta Hydrolase likely to Be Involved in the Perception of the Plant Branching Hormone, Strigolactone. Curr Biol. 22:2032–2036.

Jiang L, Liu X, Xiong G, Liu H, Chen F, Wang L, Meng X, Liu G, Yu, H, Yuan Y, et al. 2013. DWARF 53 acts as a repressor of strigolactone signalling in rice. Nature 504:401–405.

Kagiyama M., Hirano Y, Mori T., Kim SY, Kyozuka J, Seto Y., Yamaguchi S., Hakoshima T. 2013. Structures of D14 and D14L in the strigolactone and karrikin signalling pathways. Genes Cells 18:147–160.

Kato H, Ishizaki K, Kouno M, Shirakawa M, Bowman JL, Nishihama R, Kohchi T. 2015. Auxin-Mediated Transcriptional System with a Minimal Set of Components Is Critical for Morphogenesis through the Life Cycle in Marchantia polymorpha. PLoS Genet. 11:e1005084.

Kohlen W, Charnikhova T, Liu Q, Bours R. Domagalska MA, Beguerie S, Verstappen F, Leyser O, Bouwmeester H, Ruyter-Spira C. 2011. Strigolactones are transported through the xylem and play a key role in shoot architectural response to phosphate deficiency in nonarbuscular mycorrhizal host Arabidopsis. Plant Physiol. 155:974–987.

Lanfear R, Calcott B, Ho SYW, Guindon S. 2012. PartitionFinder: combined selection of partitioning schemes and substitution models for phylogenetic analyses. Molecular Biology and Evolution 29:1695–1701.

Larsson A. 2014. AliView: A fast and lightweight alignment viewer and editor for large data sets. Bioinformatics 30: 3276–3278

Lavy M, Prigge MJ, Tao S, Shain S, Kuo A, Kirchsteiger K, Estelle M. 2016. Constitutive auxin response in Physcomitrella reveals complex interactions between Aux/IAA and ARF proteins. Elife. 5:e13325.

Liang Y, Ward S, Li P, Bennett T, Leyser O. 2016. SMAX1-LIKE7 signals from the nucleus to regulate shoot development in Arabidopsis via partially EAR motif-independent mechanisms. Plant Cell. 28:1581–1601.

Lopez-Obando M, Conn CE, Hoffmann B, Bythell-Douglas R, Nelson DC, Rameau C, Bonhomme S. 2016. Structural modelling and transcriptional responses highlight a clade of PpKAI2-LIKE genes as candidate receptors for strigolactones in Physcomitrella patens. Planta 243:1441–1453.

López-Ráez JA, Charnikhova T, Gómez-Roldán V, Matusova R, Kohlen W, De Vos R, Verstappen F, Puech-Pages V, Bécard G, Mulder P, et al. 2008. Tomato strigolactones are derived from carotenoids and their biosynthesis is promoted by phosphate starvation. New Phytol. 178:863–874

Matthys C, Walton A, Struk S, Stes E, Boyer FD, Gevaert K, Goormachtig S. 2016. The Whats, the Wheres and the Hows of strigolactone action in the roots. Planta. 243:1327–1337

Miller MA, Pfeiffer W, Schwartz T editors. Proceedings of the Gateway Computing Environments Workshop (GCE). 2010 New Orleans, LA

Nakamura H, Xue YL, Miyakawa T, Hou F, Qin HM, Fukui K, Shi X, Ito E, Ito S, Park SH et al. 2013. Molecular mechanism of strigolactone perception by DWARF14. Nat Commun. 4:2613.

Nelson DC, Scaffidi A, Dun EA, Waters MT, Flematti GR, Dixon KW, Beveridge CA, Ghisalberti EL, Smith SM. 2011. F-box protein MAX2 has dual roles in karrikin and strigolactone signalling in Arabidopsis thaliana. Proc Natl Acad Sci U S A. 108:8897–8902.

Proust H, Hoffmann B, Xie X, Yoneyama K, Schaefer DG, Yoneyama K, Nogué F, Rameau C. 2011. Strigolactones regulate protonema branching and act as a quorum sensing-like signal in the moss Physcomitrella patens. Development 138:1531–1539.

Qiu, Y-L, L Li, Wang B, Chen Z, Knoop V, Groth-Malonek M, Dombrovska O, Lee J, Kent L, Rest J et al. 2006. The deepest divergences in land plants inferred from phylogenomic evidence. Proc Natl Acad Sci U S A 103:15511–15516.

Smith SM and Waters MT. 2012. Strigolactones: destruction-dependent perception? Curr Biol. 22: R924-927.

Soundappan I, Bennett T, Morffy N, Liang Y, Stanga JP, Abbas A, Leyser O, Nelson DC. 2015. SMAX1-LIKE/D53 Family Members Enable Distinct MAX2-Dependent Responses to Strigolactones and Karrikins in Arabidopsis. Plant Cell 27:3143–3159.

Stanga JP, Smith SM, Briggs WR, Nelson DC. 2013. SUPPRESSOR OF MORE AXILLARY GROWTH2 1 controls seed germination and seedling development in Arabidopsis. Plant Physiol. 163:318–330.

Stanga JP, Morffy N, Nelson DC. 2016. Functional redundancy in the control of seedling growth by the karrikin signalling pathway. Planta. 243:1397–1406.

Stirnberg P, Furner IJ, Leyser HMO. 2007. MAX2 participates in an SCF complex which acts locally at the node to suppress shoot branching. Plant J. 50:80–94.

Stirnberg P, van De Sande K, Leyser HM. 2002. MAX1 and MAX2 control shoot lateral branching in Arabidopsis. Development 129:1131–1141.

Sukumaran J, Holder MT. 2010. DendroPy: A Python library for phylogenetic computing. Bioinformatics 26: 1569–1571

Sun YK, Flematti GR, Smith SM, Waters MT. 2016. Reporter Gene-Facilitated Detection of Compounds in Arabidopsis Leaf Extracts that Activate the Karrikin Signalling Pathway. Front Plant Sci. 7:179–9

Thompson JD, Higgins DG, Gibson TJ. 1994. CLUSTAL W: improving the sensitivity of progressive multiple sequence alignment through sequence weighting, position-specific gap penalties and weight matrix choice. Nucleic Acids Res. 22:4673–4680.

Timme RE, Bachvaroff TR, Delwiche CF. 2012. Broad phylogenomic sampling and the sister lineage of land plants. PLoS One. 7:e29696.

Toh S, Holbrook-Smith D, Stogios PJ, Onopriyenko O, Lumba S, Tsuchiya Y, Savchenko A, McCourt P. 2015. Structure-function analysis identifies highly sensitive strigolactone receptors in Striga. Science. 350:203–207.

Tsuchiya Y, Yoshimura M, Sato Y, Kuwata K, Toh S, Holbrook-Smith D, Zhang H, McCourt P, Itami K, Kinoshita T et al. 2015. Probing strigolactone receptors in Striga hermonthica with fluorescence. Science. 2015 349:864–868.

Waldie T, McCulloch H, Leyser O. 2014. Strigolactones and the control of plant development: lessons from shoot branching. Plant J. 79:607–622.

Wang L, Wang B, Jiang L, Liu X, Li X, Lu Z, Meng X, Wang Y, Smith SM, Li J. (2015). Strigolactone Signalling in Arabidopsis Regulates Shoot Development by Targeting D53-Like SMXL Repressor Proteins for Ubiquitination and Degradation. Plant Cell 27:3128–3142.

Wang Y, Sun S, Zhu W, Jia K, Yang H, Wang X. 2013. Strigolactone/MAX2-induced degradation of brassinosteroid transcriptional effector BES1 regulates shoot branching. Dev Cell. 27:681–688.

Waters MT, Nelson DC, Scaffidi A, Flematti GR, Sun YK, Dixon KW, Smith SM. 2012. Specialisation within the DWARF14 protein family confers distinct responses to karrikins and strigolactones in Arabidopsis. Development 139:1285–1295.

Waters MT, Scaffidi A, Moulin SL, Sun YK, Flematti GR, Smith SM. 2015. A Selaginella moellendorffii Ortholog of KARRIKIN INSENSITIVE2 Functions in Arabidopsis Development but Cannot Mediate Responses to Karrikins or Strigolactones. Plant Cell 27:1925–1944.

Waters MT, Gutjahr C, Bennett T, Nelson D. 2017. Strigolactone signalling and evolution. Ann Rev Plant Biol 68:8–31.

Wickett NJ, Mirarab S, Nguyen N, Warnow T, Carpenter E, Matasci N, Ayyampalayam S, Barker MS, Burleigh JG, Gitzendanner MA et al. 2014. Phylotranscriptomic analysis of the origin and early diversification of land plants. Proc Natl Acad Sci U S A. 111:E4859-4868.

Wodniok S, Brinkmann H, Glöckner G, Heidel AJ, Philippe H, Melkonian M, Becker B. 2011. Origin of land plants: do conjugating green algae hold the key? BMC Evol Biol. 11:104.

Xu Y, Miyakawa T, Nakamura H, Nakamura A, Imamura Y, Asami T, Tanokura M. 2016. Structural basis of unique ligand specificity of KAI2-like protein from parasitic weed Striga hermonthica. Sci Rep. 6:31386

Yao R, Ming Z, Yan L, Li S, Wang F, Ma S, Yu C, Yang M, Chen L, Chen L, et al. 2016. DWARF14 is a non-canonical hormone receptor for strigolactone. Nature 536:469–473.

Zhao LH, Zhou XE, Wu ZS, Yi W, Xu Y, Li S, Xu TH, Liu Y, Chen RZ, Kovach A et al. 2013. Crystal structures of two phytohormone signal-transducing α/β hydrolases: Karrikin-signalling KAI2 and strigolactone-signalling DWARF14. Cell Res. 23:436–439.

Zhao LH, Zhou XE, Yi W, Wu Z, Liu Y, Kang Y, Hou L, de Waal PW, Li S, Jiang Y, et al. 2015. Destabilization of strigolactone receptor DWARF14 by binding of ligand and E3-ligase signalling effector DWARF3. Cell Res. 25:1219–1236.

Zhou F, Lin Q, Zhu L, Ren Y, Zhou K, Shabek N, Wu F, Mao H, Dong W, Gan L, et al. 2013. D14-SCF(D3)-dependent degradation of D53 regulates strigolactone signalling. Nature 504:406–410.

Zwickl DJ. 2006. Genetic algorithm approaches for the phylogenetic analysis of large biological sequence datasets under the maximum likelihood criterion. The University of Texas Austin.

